# Structural basis of archaeal RNA polymerase transcription elongation and Spt4/5 recruitment

**DOI:** 10.1101/2024.01.05.574147

**Authors:** Daniela Tarau, Felix Grünberger, Michael Pilsl, Robert Reichelt, Florian Heiß, Sabine König, Henning Urlaub, Winfried Hausner, Christoph Engel, Dina Grohmann

**Affiliations:** Institute of Microbiology & Archaea Centre, Single-Molecule Biochemistry Lab, University of Regensburg, 93053 Regensburg, Germany; Regensburg Center for Biochemistry, Structural Biochemistry Group, University of Regensburg, Regensburg, Germany; Bioanalytic Mass Spectrometry, Max Planck Institute for Multidisciplinary Sciences, 37077 Göttingen, Germany; Department of Clinical Chemistry, University Medical Center Göttingen, 37075 Göttingen, Germany Affiliation Henning; Regensburg Center of Biochemistry (RCB), University of Regensburg, 93053 Regensburg, Germany

**Keywords:** Archaea, cryo-EM, RNA polymerase, transcription elongation, transcription, Spt4/5

## Abstract

Archaeal transcription is carried out by a multi-subunit RNA polymerase (RNAP) that is highly homologous in structure and function to eukaryotic RNAP II. Among the set of basal transcription factors, only Spt5 is found in all domains of life but Spt5 has been shaped during evolution, which is also reflected in the heterodimerization of Spt5 with Spt4 in Archaea and Eukaryotes. To unravel the mechanistic basis of Spt4/5 function in Archaea, we performed structure-function analyses using the archaeal transcriptional machinery of *Pyrococcus furiosus* (*Pfu*). We report single-particle cryo-electron microscopy reconstructions of apo RNAP and the archaeal elongation complex (EC) in the absence and presence of Spt4/5. Surprisingly, *Pfu* Spt4/5 also binds the RNAP in the absence of nucleic acids in a distinct super-contracted conformation. We show that the RNAP clamp/stalk module exhibits conformational flexibility in the apo state of RNAP and that the enzyme contracts upon EC formation or Spt4/5 engagement. We furthermore identified a contact of the Spt5-NGN domain with the DNA duplex that stabilizes the upstream boundary of the transcription bubble and impacts Spt4/5 activity *in vitro*. This study, therefore, provides the structural basis for Spt4/5 function in archaeal transcription and reveals a potential role beyond the well-described support of elongation.

## Introduction

During the first step in gene expression, genomic DNA is transcribed in a highly regulated manner to synthesize the cellular RNA pool. The cyclic and multistep transcription process can be divided into three main steps: initiation, elongation, and termination^1–5^. Although Archaea are prokaryotes, the components of the archaeal transcription machinery do not resemble the bacterial transcription apparatus but are highly homologous to the eukaryotic RNAP II transcription machinery^3,4^. This applies to the composition and structure of the RNAP itself and extends to the basal transcription initiation and elongation factors. Eukaryotic Pol I and III even retain structural elements lost in Pol II, suggesting a common archaeal origin^6–8^. In Archaea, transcription initiation depends on the TATA-binding protein (TBP) and transcription factor B (TFB), which recognize the promoter DNA forming sequence-specific interactions with the TATA-box and B-recognition element (BRE), respectively^9–15^. The helicase-independent melting of the promoter DNA around the transcription start site is supported by transcription factor E (TFE), leading to the efficient formation of the open pre-initiation complex(PIC)^10,16–19^. Productive RNA synthesis becomes possible when contacts between initiation factors and RNAP are broken, and RNAP escapes from the promoter.

Transition into the elongation phase is accompanied by the replacement of TFE by the elongation factor Spt4/5^11,12,17^. Spt5 is the only conserved transcription factor found in all domains of life^3^. Like its bacterial counterpart NusG, archaeal Spt5 comprises an N-terminal NusG domain (NGN), a linker domain, and a C-terminal Kypridis-Ouzounis-Woese (KOW) domain^20^. Archaeal and eukaryotic Spt5 form stable heterodimers with the small protein Spt4^20–23^. In Eukaryotes, however, Spt5 is much larger and includes four to seven consecutive KOW domains at its C-terminus. Productive RNA synthesis during the elongation phase is not continuous but is frequently interrupted by pausing events. Upon interaction with the elongating polymerase, Spt4/5 reduces pausing, thereby increasing the processivity of RNAP^24^. Moreover, Spt4/5 promotes transcription through chromatinized DNA templates in Archaea and Eukaryotes^20,25^. In Bacteria, NusG suppresses pausing and backtracking of the RNAP and interacts with effector proteins such as the termination factor Rho via its KOW domain^26,27^.Furthermore, it was shown that NusG mediates the coupling between RNAP and the ribosome contacting the RNAP clamp and the ribosomal subunit S10^28–31^. In contrast to NusG in *E. coli*, a gene-specific homologue called RfaH and *B. subtilis* ‘NusG-like’ homologues interact with the non-template strand of the transcription bubble and thereby enhance transcriptional pausing^32–35^.

Structural, single-molecule and biochemical data revealed the mechanistic basis for the stimulatory effect of NusG and Spt4/5. Spt5, as well as NusG, binds to the tip of the RNAP clamp coiled-coil motif via a hydrophobic cavity in the NGN domain and bridge over the DNA binding channel to the RNAP lobe domain^21,22^. The clamp is a domain mostly conserved in structure among multi-subunit RNAPs and can adopt open or closed conformations. While the degree of clamp contraction varies among transcription systems, especially by the eukaryotic RNAPs, a closed clamp defines actively elongating RNAPs. At the same time, an expanded or partially expanded conformation is found in inactive enzymes^36^. NusG prevents conformational changes that are connected to pausing states, e.g. the opening of the clamp^35^. Archaeal Spt4/5 induces the closure of the RNAP clamp when the initiation complex transitions to the elongation complex, thereby securing the DNA template in the DNA binding channel rendering the elongation complex more stable^19^. Furthermore, Spt4/5 and RNAP subunits Rpo4/7 are required for efficient termination mediated by the extrinsic termination factor aCPSF1 in Archaea, suggesting that both protein complexes provide interaction sites for aCPSF1^37,38^.

High-resolution crystal structures and structural models of bacterial and eukaryotic transcription complexes derived from cryo-electron microscopy (cryo-EM) and single particle analysis data provided insights into the molecular basis of transcription initiation, elongation, transcriptional pausing, and termination^1,5,39–44^. However, structural information on archaeal transcription complexes is sparse and only includes structures of the RNAP apo enzyme^45–49^, of RNAP in complex with DNA^50^ or TFE, as well as a complex formed of RNAP, DNA and TFE^48^. Furthermore, structures of RNAP in complex with TFS4^51^or an archaeon-viral inhibitor^49^were published. Additionally, an isolated clamp fragment in complex with archaeal Spt4/5^21^ was structurally described, and a low-resolution structural model of the archaeal RNAP-Spt4/5 complex in the absence of nucleic acids is available, but structural details could not be discerned^22^. Distance constraints derived from a comprehensive single-molecule FRET study were used to build a structural model of the full archaeal PIC^16^.

To gain insights into the structural (re-)organization of archaeal transcription elongation complexes (TEC) in the absence and presence of Spt4/5 and to understand the mechanisms of Spt4/5 action, we determined single-particle cryo-EM reconstructions of the archaeal RNAP apo enzyme from *Pyrococcus furiosus* with and without Spt4/5 and the archaeal TEC with and without Spt4/5. Combined with biochemical and mass spectrometry data, we found that a subfraction of the cellular apo RNAP pool forms a complex with Spt4/5 in the absence of DNA or RNA. In the context of the TEC, Spt4/5 occupies a slightly different position relative to the RNAP compared to the apo RNAP-Spt4/5 complex. Furthermore, we found that archaeal Spt4/5 interacts with the upstream DNA representing an organism-specific interaction mode. Transcription assays with Spt5 mutants revealed that this interaction contributes to the processivity of the archaeal RNAP. Our data, therefore, add to the structural and mechanistic understanding of archaeal transcription complexes and the functional role of Spt5.

## Material and Methods

### Synthetic oligonucleotides

Elongation complexes were formed using an elongation scaffold composed of a fully complementary template and non-template strand and a short (14 nt) or slightly longer (22 nt) RNA oligonucleotide complementary to CGCCTGGTC (9) bases of the template strand. All DNA and RNA oligonucleotides were synthesized by Microsynth AG. Sequences are as follows: template strand (TS) 5’- CCACCCTTACCTCCACCATATGGGAGATCCATTAGAGTAGCGCCTGGTCATTACTTAAGATGAAGTAGTTATAGTACTGCCGG-3’; non-template strand (NTS) 5’- CCGGCAGTACTATAACTACTTCATCTTAAGTAATGACCAGGCGCTACTCTAATGGATCTCCCATATGGTGGAGGTAAGGGTGG-3’; partially complementary RNA (short) 5‘-AUUUAGACCAGGCG-3‘; and long RNA 5‘- UUUUUUUUAUUUAGACCAGGCG-3’. The oligonucleotides used for the genomic modification are listed in Table S2.

### Genomic modification of *rpoD* and cultivation of *Pyrococcus furiosus*

To enable a simplified purification of the cellular RNAP we introduced a StrepII-His6-tag at the C-terminal end of subunit Rpo3 using an established genetic system for *P. furiosus* DSMZ 3638^78,79^. We used double crossover recombination with plasmid pMUR1 to modify the genome. This plasmid contains *rpoD* (PF1647) together with 250 bp of the upstream region. The corresponding sequence was amplified from wild-type DNA with the primer pair 1648/1647-BH1-F and PF1647_Fus-StrepII_Ri. The StrepII-His6-tag at the C-terminal end was provided by a double-stranded oligonucleotide (primer Fus_Strep_His_F and Fus_Strep_His_R). For the selection of the *Pyrococcus* transformants with the antibiotic simvastatin, the coding sequence of the hydroxymethylglutaryl CoA reductase from *Thermococcus kodakarensis*, was used. The corresponding sequence was amplified from plasmid pMUR27 using the primer pair PF1647upTk0914F and TK0914-R^79^. Furthermore, the plasmid includes a 950 bp sequence downstream of *rpoD* to enable double crossover recombination. This fragment was amplified from genomic DNA using the primer pair Tk0914PF1647dwF and Fus004_BamHI-R. All the DNA fragments were combined step-by-step by single overlap extension PCR. The fused fragment was hydrolyzed with BamHI und ligated into a pUC19 vector, which was also hydrolyzed with BamHI and dephosphorylated with calf intestinal phosphatase.

The resulting plasmid pMUR1 was sequenced and 1 µg of the linearized version was used for transformation of *Pfu* as described previously^78,79^. Transformants were enriched with 10 µM simvastatin in ½ SME-starch liquid medium at 85°C for 48 h, and pure cultures were isolated after plating the cells on solidified medium in the presence of 10 µM simvastatin. The integration of the DNA fragment into the genome via double crossover was verified by sequencing of corresponding PCR products.

*Pyrococcus furiosus* cells were first grown for 6 hours in a volume of 20 ml under anaerobic conditions in ½ SME medium supplemented with 0.1% (w/v) yeast extract and 0.05% (w/v) Na_2_S according to previously established protocols^80^. The preculture was incubated at 90°C with N_2_ gas phase at 200 kPa pressure until a cell density of 7.5 x 10^6^ cells was reached. The preculture was used to inoculate a 100L fermenter with ½ SME medium supplemented with 0.1% peptone, 0.1% yeast extract, 0.1% starch and 0.05% (w/v) Na_2_S. The pH was adjusted with 5M NaOH to 7.5 and cells were incubated for 12 hours at 90°C and stirred at 250 rpm. The gas phase was N_2_/CO_2_ (80%:20% ratio, respectively) and 200 kPa pressure. 117 grams of pellet were obtained after harvesting by centrifugation at 20144 g, with Centrifuge 17S/RS (Heraeus Sepatech) with a continuous flow rotor 3049 (titanium). The pellet was split into 3 aliquots, plunge-frozen and stored at -80°C until further use.

### Purification of RNA polymerase from *Pyrococcus furiosus*

The cell pellet was resuspended in 2.5 ml per g of cells lysis buffer (100 mM Tris/HCl pH 8.0, 1 M NaCl, 20 mM imidazole, 20% (v/v) glycerol, 2.5 mM MgCl_2_) 1 EDTA free protease inhibitor cocktail tablet (Roche) per 50 mL of lysis buffer. 0.5 g of glass beads (0.106 µm diameter, Sigma-Aldrich) were added to the mixture and the pellet was resuspended by bead beating using the FastPrep-24™ 5G bead beating grinder and lysis system from MP at a speed of 6.5 m/sec for 40 seconds. Cells were lysed by sonication using Bandelin Sonopuls HD 2070 at 60% amplitude for 6 minutes. Sonication was repeated five to six times. Cell lysis was confirmed by light microscopy. After lysis, the crude extract was centrifuged at 15,000 g for 10 minutes (Eppendorf 5810 centrifuge) to pellet cell debris and glass beads. The supernatant was transferred and subjected to ultracentrifugation at 100,000 g using a type 70 Ti Rotor in the Optima Le-80K Ultracentrifuge (Beckmann).

The supernatant was carefully collected and filtered using 0.45 µm syringe filters (Cytiva). The polymerase was purified using affinity chromatography exploiting the 6xHis-tag. To this end, the filtrate was applied to a HisTrap™ 5 ml Fast Flow (Cytiva) equilibrated in lysis buffer using an Äkta Purifier system (GE Healthcare). The column was washed with lysis buffer (100 mM Tris/HCl pH 8.0, 1 M NaCl, 20 mM imidazole, 20% (v/v) glycerol, 2.5 mM MgCl_2_) until the absorbance at 280 nm and 260 nm dropped close to 0 mAU. The content was eluted from the column using 30 ml elution buffer (100 mM Tris/HCl pH 8.0, 300 mM NaCl, 250 mM Imidazole, 20% (v/v) glycerol, 2.5 mM MgCl_2_). Subsequently, the RNA polymerase was further purified performing a size exclusion chromatography step (column: HiLoad 26/600 Superdex 200 PG from Sigma-Aldrich). The following running buffer was used for the size exclusion chromatography: 100 mM HEPES/NaOH pH 8.0, 150 mM NaCl, 20% (v/v) glycerol, 2.5 mM MgCl_2_. Elution fractions were loaded on a denaturing SDS polyacrylamide gel (PAGE) and fractions containing the RNA polymerase were aliquoted and stored at -80°C until further use. The concentration of 1.638 mg/ml was determined by absorption spectroscopy at 280 nm wavelength using a NanoDrop™ One/OneC Microvolume UV-Vis Spectrophotometer (Thermofisher).

### Recombinant Spt4/5 expression and purification

The genes encoding Spt4 (GeneID 41712045) and Spt5 (GeneID 41713813) were amplified by PCR from *Pfu* genomic DNA and further were cloned into pET14b and pET17b expression vectors, respectively by Gibson Assemby^81^. The *spt4* gene was genetically fused to a sequence that encodes a 6x histidine tag at the C-terminus of Spt4, whereas Spt5 was cloned without any additional tag. 200 µg of each plasmid were used to transform *E. coli* Rosetta™(DE3) competent cells (Merk, Darmstadt) using heat transformation. The cell suspension was plated on LB-agar plates containing 1 µg/ml ampicillin. Plates were incubated at 37°C overnight. The next day, one positive colony from each plate was resuspended in 20 µl LB medium and plated again on a LB agar plate that contained 1 µg/ml ampicillin and the plate was incubated again at 37°C overnight. The next day, the bacterial colonies were resuspended in 5 ml LB liquid medium and this solution was used to inoculate a 1L culture in LB medium using a starting OD_600_ of ∼0.07. The cultures were incubated at 37°C shaking at 160 rpm. Protein expression was induced by addition of Isopropyl-beta-D-Thiogalactosid (IPTG) to a final concentration of 1 µg/ml at an OD_600_ 0.6-0.7. After the induction, the cells were further incubated for 4 hours at 37°C shaking at 160 rpm. Cells were harvested by centrifugation for 10 min at 5500 g at 4°C. The pellets were resuspended in low salt buffer (40 mM HEPES/KOH pH 7.4, 50 mM NaCl, 15% (v/v) glycerol) and sonicated five times for 30 seconds using a Bandelin Sonopuls HD 2070 at 50% amplitude and cycle 5. Cell debris was eliminated by centrifugation (JA-10 fixed rotor; 1 hour; 20,000 g; 4°C). Spt4 and Spt5 maintain their thermostability even when recombinantly expressed, therefore the crude extract was incubated at 80°C for 20 minutes to heat denature and precipitate most of the *E. coli* proteins. The precipitates were pelleted by centrifugation (JA-25.50 fixed rotor; 20.000 g; 15 minutes; 4°C) and the supernatant was filtered using a 0.2 µm syringe filter (Cytiva). At this point, the lysates that contained either Spt4 or Spt5 were mixed. The Spt4/5 complex was purified via affinity chromatography employing the His-tag of Spt4. Affinity purification was performed using the Äkta Purifier system and a HisTrap™ 1 ml Fast Flow (Cytiva) column equilibrated in low salt buffer. The lysate was loaded onto the column and washed in low salt buffer until the absorbance at 280 nm dropped close to 0 mAU. Spt4/5 was eluted using a gradient elution (elution buffer: 40 mM HEPES/KOH pH 7.4, 100 mM NaCl, 15% (v/v) glycerol, 500 mM imidazole). Elution fractions containing Spt4/5 were identified using SDS PAGE analysis and subjected to size exclusion chromatography (Superdex® 75 10/300 GL; Sigma-Aldrich). The size exclusion run was performed using isocratic elution in low salt buffer (40 mM HEPES/KOH, pH 7.4, 50 mM NaCl, 15% glycerol (v/v)). The purification success was assessed by SDS-PAGE. The concentration was determined by using an extinction coefficient of 15,930 calculated with ProtParam tool (Expasy)^82^ and with absorbance at 280 nm using NanoDrop™ One/OneC Microvolume UV-Vis Spectrophotometer (Thermofisher).

#### Spt5 mutagenesis

Residues H12, R64, H65 and R67 were mutated to alanines via mutagenic primer PCR (Table S2). The forward mutagenesis primer was phosphorylated at the 5’ end and contained the mutated sequence at the respective position. For what concerns the reverse primer, it is designed to cover the complementary sequence just upstream of the forward primer. After gradient PCR the successful amplification was verified by 1% agarose gel electrophoresis. 5 µl from the PCR product were circularized using 1 µl T4 DNA ligase and 2 µl ligation buffer 10x (NEB) in a final volume of 20 µl. To stop the ligation reaction, the sample was incubated at 60°C for 1 hour to denature the T4 DNA ligase. 5 μl from the ligation mixture was used to transform 100 μl *E. coli* DH5α competent cells. Transformants were selected on 100 μg/ml ampicillin-containing LB agar plates (incubation of plates overnight at 37°C). A colony was collected and resuspended in 5 ml liquid LB medium with 100 μg/ml ampicillin and cultivated for overnight at 37°C. Plasmid isolation was performed using the Plasmid Miniprep Kit, peqGOLD Plasmid Miniprep Kit I (VWR Life Science). Successful mutagenesis was verified by Sanger sequencing.

#### Spt5 mutant purification

Spt5 mutants were overexpressed as described for the wild-type protein. Heterodimer formation with Spt4 was performed on a HisTrap™ 1 ml Fast Flow (Cytiva) to which 1 mg of purified Spt4-6xHis protein was immobilized. Bacterial lysate containing a Spt5 mutant was applied over the column. The Spt4/5 complex was eluted using a gradient elution (elution buffer consisted of 40 mM HEPES/KOH pH 7.4, 100 mM NaCl, 15% v/v glycerol, 500 mM imidazole). The Spt4/5 heterodimer was further purified by size exclusion chromatography (Superdex 75 10/300 GL, GE Healthcare) through isocratic elution in low salt buffer. Fractions containing Spt4/5 were evaluated and identified by SDS-PAGE, 16% v/v polyacrylamide concentration.

### Elongation complex assembly for cryo electron microscopy analysis

Transcription elongation complexes (TEC) were assembled by first annealing the RNA (400 µM) to the TS (400 µM) using a molar ratio of 9:1 in a final volume of 40 µl and incubating this sample at 95°C for 3 minutes and slow cooling. In a second step, RNAP is added to the pre-annealed RNA-TS hybrid (40 µM) using a 1:3 molar ratio by incubation at 80°C for 15 minutes. In a last step the NTS is added to the complex with 3 times excess in molarity with respect to the RNAP by incubation at 80°C for 10 minutes. The final volume of the reaction was 100 µl.

For the assembly of the TEC in absence of Spt4/5 the short RNA was used for the elongation scaffold, whereas for the TEC with Spt4/5 sample, the long RNA oligonucleotide was used instead.

To increase the occupancy of Spt4/5 in the TEC, a BS^3^ crosslinker (Thermo Fisher Scientific) was added to the preformed complex at a final concentration of 3 mM. TEC and Spt4/5 were incubated at 30°C for 30 minutes. The reaction was quenched by incubation at 30°C for 30 minutes with Lys/Asp mixture (0.1 mM) and incubation at 20°C for 20 minutes with ammonium bicarbonate (0.02 mM).

The excess of crosslinker, quencher and precipitates were eliminated by gel filtration chromatography using the same column and buffer as described for the purification for Spt4/5 wt. The crosslinking efficiency was checked using a 4% to 20% gradient SDS-PAGE.

### Cryo-EM grid preparation, data acquisition and processing

The overall workflow described in Pilsl et al.^83^was adapted to *Pfu* RNA polymerase complexes. Apo RNA polymerase. 100 µg of apo RNAP were run onto Superose 6 Increase 3.2/300 column in cryo-buffer (100 mM HEPES/KOH pH 8, 100 mM NaCl, 2.5 mM MgCl_2_). The 50 µl elution fractions corresponding to the polymerase peak were collected and used for vitrification. The sample was applied onto freshly glow discharged with EasiGlow, (TedPella) (1x 0.4 mbar; 15 mA; 30seconds) UltrAufoil R1.2/1.3, 300 mesh gold grids (Quantifoil) and plunge-frozen into liquid ethane (Vitrobot Mark IV; Thermo Fischer Scientific; 100% humidity; 4°C; 3 seconds blotting time, 0 blotting force). Data collection was performed at a CRYO ARM™ 200 (Jeol) equipped with Gatan K2 summit. A total of 8,212 movies in super resolution mode at 50,000x magnification and a pixel size of 0.968 Å/pixel, the in-column energy filter was operated with a slit width of 20 eV. A total exposure of 40 e-/A^2^ was fractionated over 40 frames (1 e-/pixel/s) at a defocus range from -0.6 µm to -2 µm using SerialEM software^84^.

Transcription elongation complex. 100 µg of RNAP were used to assemble a TEC on a synthetic elongation scaffold as described in the section “Elongation complex assembly for cryo electron microscopy analysis”. TECs were applied to a Superose 6 Increase 3.2/300 column and run in cryo-buffer (100 mM HEPES/KOH pH 8, 100 mM NaCl, 2.5 mM MgCl_2_). The fractions corresponding to the polymerase peak were collected and used for vitrification. The sample was applied onto freshly glow discharged with EasiGlow, (TedPella) (2x100 sec; 0.4 mbar; 15 mA) UltrAufoil R1.2/1.3 gold grids, 300 mesh (Quantifoil) and plunge-frozen into liquid ethane (Vitrobot Mark IV; Thermo Fischer Scientific; 100% humidity; 4°C; 3 seconds blotting time, 15 blotting force). Data was acquired on a Titan Krios G3 Electron microscope equipped with a Falcon III detector using EPU software (Thermo Fisher Scientific). Movies were recorded at 75,000x magnification (pixel size 1.0635 Å/pixel) in linear mode at a defocus ranging from -1.2 µm to -2.0 μm. A total exposure of 86 e–/Å^2^ was fractionated over 40 frames, the electron dose was 19 e-/pixel/s.

Transcription Elongation Complex with Spt4/5. 100 µg of RNAP were used to assemble on the synthetic scaffold and run onto Superose 6 Increase 3.2/300 column in cryo-buffer (100 mM HEPES/KOH pH 8, 100 mM NaCl, 2.5 mM MgCl_2_). 50 µg from the most concentrated fractions were collected and used to crosslink the TEC to Spt4/5 using a BS^3^ crosslinker (description of the crosslinking procedure is outlined in the section “Elongation complex assembly for cryo electron microscopy analysis”). BS^3^-crosslinking can stabilize the association for transcription factors, without influencing the architecture of their complexes^85^. Elution fractions from a second size exclusion chromatography run corresponding to the polymerase peak were collected and used for vitrification. The sample was applied onto freshly glow discharged with EasiGlow, (TedPella) (2x 100 sec, 0.4 mbar; 15 mA) UltrAufoil R1.2/1.3, 300 mesh gold grids (Quantifoil) and plunge-frozen into liquid ethane (Vitrobot Mark IV; Thermo Fischer Scientific; 100% humidity; 4°C; 3 seconds blotting time, 15 blotting force). Data collection was performed at a CRYO ARM™ 200 (Jeol) equipped with Gatan K2 summit in super resolution mode using SerialEM software^84^. Movies of 40 frames were acquired at 50,000x magnification (pixel size 0.968 Å). The electron dose was 1 e– /px/s with a total of ∼40 e–/Å^2^. The defocus span was of -0.6 μm till -2 μm with 0.2 μm steps.

### Data processing and model building

Apo RNA polymerase. 8,212 movies of 40 frames each were collected and processed in RELION 4.0 (processing tree **Figure S3**, **Supplementary Figure S4**)^86^. The movie frames were aligned, and dose weighted using RELION’s implementation MotionCor2^87^ and Contrast Transfer Function (CTF) estimation using GCTF. 6,482 micrographs were selected for further evaluation based on astigmatism values, ice contamination or defocus values. Particles were first picked using Laplacian-of-Gaussian algorithm and an initial 2D classification was run. The best classes were used for Topaz^88^ training and picking. 2D classification was performed twice and the best classes were selected for unbiased 3D reconstruction using 3D Initial Model function in RELION with 4 classes. After sequential 3D classification and filtering unaligned particles or contaminants, 2 classes corresponding to 2 different conformations: open clamp conformation and closed clamp conformation with Spt4/5, were processed further separately. 196,232 particles for the open conformation were CTF refined and Polished with Bayesian Polishing function leading to 3.5 Å map which was further classified without image alignment, using a mask focused onto the clamp and stalk domain. 161,531 particles corresponding to the best aligned classes were further refined yielding to a 3.2 Å resolution map.

For what concerns the closed conformation with Spt4/5, a mask incorporating the clamp extra-density was created and 3D classification without image alignment was performed. Particles corresponding to the best classes were selected, CTF corrected and polished. A final map of 3.4 Å was obtained. Final resolution was calculated using a cut-off at 0.143 gold standard in 3DFSC^89^ online tool. To better describe the conformational heterogeneity of the polymerase, we re-analysed the dataset using cryoSPARC^52^. Contamination and badly aligned particles were removed in two rounds of 2D classification. Particles were refined on a newly generated initial model and 3D classification separated Spt4/5 bound und unbound polymerase. Both sets of particles were subsequently refined and the conformational variability was evaluated by 3D variability analysis (3DVA). For the unbound RNAP, we firstly sorted out putative dimeric RNAPs, using 2D and 3D classification, to avoid bias of multimeric enzyme conformation. Principal component analysis indicated flexibility in clamp domain and polymerase contraction (see **Figure S4** and **Supplementary Movie S3**). Particles were sorted into clusters along one component, a subset of 111,194 particles was selected and refined, resulting in a 3.0 Å reconstruction of the contracted apo enzyme (in closed clamp conformation, **Figure 1b**).

**Figure 1:**
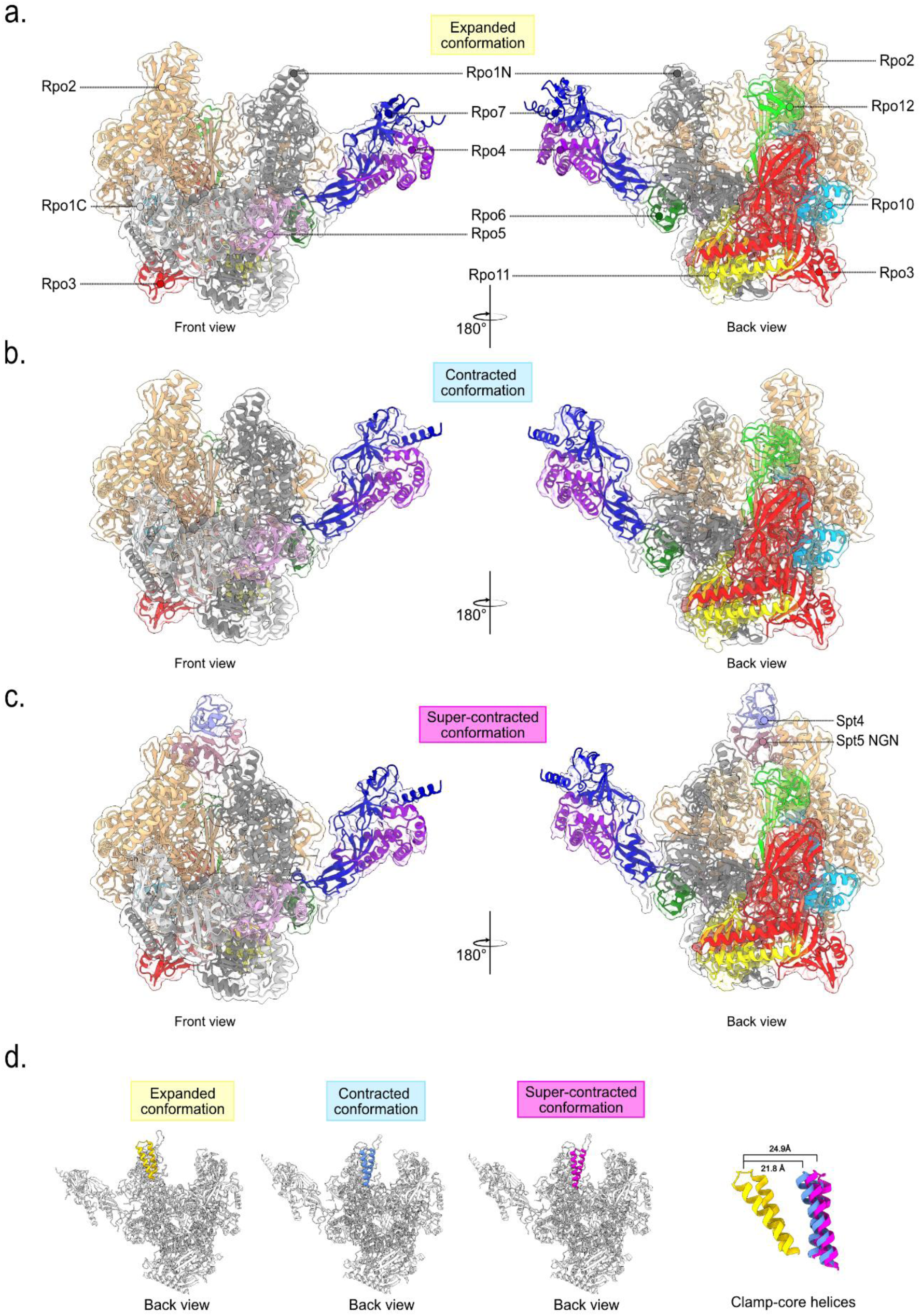
Structure of the euryarchaeal RNA polymerase apo enzyme and RNA polymerase-Spt4/5 complex from *Pyrococcus furiosus*. **(a)** Structural model of the apo RNA polymerase from *Pfu* with an open clamp in two different views with the corresponding subunit denomination. **(b)**Cryo-EM structure of *Pfu* apo RNA polymerase in a closed clamp conformation in two different views. **(c)** Cryo-EM reconstruction of the RNA polymerase with Spt4/5 shows that the clamp domain adopts a closed conformation. **(d)** Direct comparison of *Pfu*RNAP structures with open and closed clamp and close-up view of the clamp coiled coil helices that undergo a movement of 21.4 Å when upon transition from the open to closed state. Structures were globally aligned in ChimeraX, contraction distance was measured between the Cα positions of Proline 261within subunit Rpo1N in the three models.

3DVA of the Spt4/5 bound polymerase indicated flexibility in the jaw and lobe domain, inducing polymerase contraction (**Supplementary Movie S4**). The lobe domain appears to contact Spt5 and might stabilize its association. We could not sort a distinct state, EM density in this part was still poor and we refrain from modelling this interaction.

Transcription Elongation Complex. The data collected was 2,952 movies with 40 frames each processed in RELION 3.1 and RELION 4.0 beta (processing tree FigureS8). The frames were aligned, and dose weighted as described before. 1,229,339 particles picked with crYOLO^90^ were classified in 2D. Around 90% of the starting particles were used for generating 3D model using Initial model option on RELION. Sequential 3D classifications were performed to separate best aligned particles from contamination. Finally, 476,655 particles were CTF refined and polished to a final resolution of 3.1 Å determined with 3DFSC online tool at 0.143 cut-off gold standard.

Transcription Elongation Complex bound to Spt4/5. 6076 movies of 40 were corrected as described before. Particles were first picked with Laplacian-of-Gaussian algorithm to generate 2D classes to be used for a second round of particle picking using reference-based picking algorithm (processing tree Figure S10). Picked particles were extracted with a binning factor of 4x and 2D classification was done. A first 3D model was generated with the Initial model function in RELION, followed by sequential 3D classification and finally 322,138 particles were converted to the original pixel size. CTF correction and Bayesian polishing was performed followed by another round of CTF refinement. A 3.3 Å resolution map was generated which was further classified without image alignment with a mask focused to the clamp and stalk domain eliminating particles not presenting Spt4/5 density on the clamp domain. The selected classes were classified again without image alignment enlarging sightly the mask. Particles corresponding to the best class were refined yielding to 3.45 Å resolution map determined as previously. To improve local sharpening for easier model interpretation, LocScale^91^ software from CCP-EM^92^ was used on this map.

Model building was performed on the TEC map with a homology model created on SwissModel^93^ (Expasy) using *Themooccus kodakarensis* RNA polymerase (PDB: 6kf9) as template model. The homology model was then fitted into the map using WinCoot 0.9.8.1^94^ and Isolde^95^ from ChimeraX 1.5^96^. The models were refined in PHENIX 1.20.1-4487^97^.

### Sample preparation for mass spectrometry analysis

*Pyrococcus furiosus* pMUR1 and HisOnly strain were resuspended from a fresh capillary from the Regensburg Bacterial Bank and cultivated in 40 ml ½ SME medium supplemented with 0.1% (w/v) yeast extract, 0.1% (w/v) peptone, 0.4% (w/v) sodium pyruvate and 0.05% (w/v) Na_2_S and using a N_2_ gas phase (80kPa pressure). The medium was furthermore supplemented with 10 µM simvastatin to restrict the growth to the *Pfu* mutant that carries the genetically modified *rpoD* locus. The “His-tag only” mutant was a genetically engineered strain with the strategy described in Waege et al., 2010^78^ and was used as negative control since it is containing a plasmid able to express only the 6xHis-tag. The re-cultivated cultures were grown at 85°C for approximately 72 hours. Five cultures of each mutant were inoculated in the same medium in the absence of simvastatin and grown at 95°C for 12 hours. The cultures were pelleted at 4,000 g for 30 minutes at 4°C. The cells were plunge-frozen in liquid nitrogen and kept at -80°C until further use. The cells were resuspended in 700 µl homogenizing buffer (100 mM HEPES/KOH pH 8.0, 600 mM NaCl, 20% (v/v) glycerol, 20 mM imidazole, 2.5 MgCl_2_). The cells were lysed by sonication using Bandelin Sonopuls HD 2070 at 25% amplitude for three minutes. The sonication was repeated six times. Cell debris was removed by ultracentrifugation at 20,000 g for 1 hour at 4°C using a Beckman Optima MAX Ultracentrifuge and a TLA-55 rotor. The supernatant was carefully collected and loaded onto 30 µl MagneHis™ beads (Protein Purification System, Promega) and incubated for 1 hour, gently rotating. Subsequently, the supernatant was removed from the beads. Four washing steps were performed (three minutes each) using the homogenizing buffer. Finally, the beads were incubated for 15 min with 50 µl elution buffer (100 mM HEPES/KOH pH 8.0, 600 mM NaCl, 20% v/v glycerol, 500 mM imidazole, 2.5 MgCl_2_) to elute the polymerase and all interacting protein partners. One preparation was used as quality control as shown in the 4-20% gradient SDS-PAGE in (FIGURE S5). The four replicates were precipitated in 250 µl 99.5% acetone (acetone was pre-cooled to -20°C) by incubating the samples at -20°C for one hour. The precipitated protein mixture was centrifuged at 18,000 g for one hour at 4°C. The supernatant was carefully removed and the pellet was resuspended in 5 µl of 6x protein loading buffer solution (350 mM Tris/HCl pH 6.8, 30% glycerol (v/v), 10% sodium dodecyl sulphate (w/v), 0.6 M DTT, 0.012% bromophenol blue (w/v)).

### Mass spectrometry analyses

#### Sample preparation

Samples were reconstituted in 1× NuPAGE LDS Sample Buffer (Invitrogen), separated on 4-12 % NuPAGENovex Bis-Tris Minigels (Invitrogen) and stained with Coomassie Blue. Lanes were cut into 23 equidistant slices irrespective of staining. After washing, gel slices were reduced with dithiothreitol (DTT), alkylated with 2-iodoacetamide and digested with Endopeptidase Trypsin (sequencing grade, Promega) overnight. The resulting peptide mixtures were then extracted, dried in a SpeedVac, reconstituted in 2% acetonitrile/0.1% formic acid/ (v:v) and prepared for nanoLC-MS/MS as described previously (Atanassov and Urlaub, 2013^98^). nanoLC-MS/MS analysis for mass spectrometric analysis samples were enriched on a self-packed reversed phase-C18 precolumn (0.15 mm ID x 20 mm, Reprosil-Pur120 C18-AQ 5 µm, Dr. Maisch, Ammerbuch-Entringen, Germany) and separated on an analytical reversed phase-C18 column (0.075 mm ID x 200 mm, Reprosil-Pur 120 C18-AQ, 1.9 µm, Dr. Maisch) using a 46 min linear gradient of 4-35 % acetonitrile/0.1% formic acid (v:v) at 300 nl min-1). The eluent was analyzed on Q ExactiveHF hybrid quadrupole/orbitrap mass spectrometer (ThermoFisher Scientific, Dreieich, Germany) equipped with a FlexIon nanospray source and operated under Xcalibur 2.5 software using Top30 data-dependent acquisition methods, respectively. One full MS scan across the 350-1600 m/z range was acquired at a resolution setting of 120,000, and AGC target of 1*10e6 and a maximum fill time of 60 ms. Precursors of charge states 2 to 6 above a 4*20e4 intensity threshold were then sequentially isolated at 1.6 FWHM isolation window, fragmented with nitrogen at a normalized collision energy setting of 30%, and the resulting product ion spectra recorded at a resolution setting of 15,000 FWHM, AGC targets of 1*10e5 and a maximum fill time of 60 ms. Selected precursor m/z values were then excluded for the following 25 s.

#### Data processing

Raw data were processed using Mascot 2.3.02 (Matrixscience, London, UK) using standard search parameters (Trypsin/P, maximum 2 missed cleavages, fixed modification C-carbamidomethyl, variable modification protein N-terminal acetylation and M-oxidation, 20 ppm precursor and fragment tolerance, 1% FDR as established by a target/decoy search against a forward/reverse concatenated database). Data were searched against the UniProtKB*Pyrococcus furious* database (7054 entries). Search results were then combined and analysed for normalized spectral counting quantification in Scaffold 5.2.1 software (Proteome Software, London, UK)

### Mass spectrometry data analysis

Preprocessing was performed to address missing values and normalize spectral counts. Briefly, zeros in the dataset were replaced with NA to accurately represent undetected proteins and log2 transformation with subsequent median normalization was applied to mitigate the mean-variance relationship. Principal component analysis (PCA) was employed to identify similarities between samples and to detect outliers, leading to the removal of outliers from the analysis (specifically, replicate 4). The proDA package (https://github.com/const-ae/proDA, v. 1.10.0) was utilized to analyze label-free mass spectrometry data. The normalized data were fitted with a linear probabilistic dropout model within the proDA framework. Experimental design, including contrasts between different groups or conditions, was explicitly specified during the analysis. To identify proteins with coefficients significantly different from zero, hypothesis testing was conducted using proDA. A one-sided Wald test was employed to detect differentially abundant proteins, with multiple hypothesis testing correction using Benjamini-Hochberg (Supplementary Tables 3 and 4).

### Immunodetection

*Pfu*RNAP, Spt4/5 wild-type, Spt4/5 mutant and Spt4 and Spt5 proteins were loaded onto a 16% SDS-polyacrylamide gel and ran at 180V constant for 40 minutes in the Bio Rad system. The proteins were transferred onto Immobilon®-PSQ PVDF Membrane (Merk) using transfer buffer (25 mM TRIS, pH 8.3, 192 mM glycine, 10% methanol (v/v)). The transfer was carried for 1 hour at room temperature 90V/350mA with constant voltage. After the transfer, the western blot sandwich was disassembled and the membrane was blocked in blocking solution (TRIS 10 mM, pH 8, NaCl 150 mM, 0.1 % Tween 20 (v/v), 5% milk powder (w/v)) and incubated with rabbit anti-Spt5 primary antibody (1:1000 volume ratio) overnight at 4°C shaking. The day after, the membrane was washed in TBST buffer (TRIS 10 mM, pH 8, NaCl 150 mM, 0.1 % Tween 20 (v/v)) and incubated in blocking solution and chicken anti-rabbit IgG (H+L) secondary antibody, Alexa Fluor™ 647 (Invitrogen) in a 1:10000 volume ratio. Alexa Fluor 647 fluorescence signal was detected at the ChemiDoc™MP Gel Imaging System (BIORAD).

### *In vitro* transcription

The TEC was assembled using Cy5-labelled RNA as part of the elongation scaffold. The fluorophore was attached to the 5’-end of the 22 nt long RNA. RNA (100 µM) and the TS (400 µM) were annealed at a ratio of 3:1 molar ratio Incubated at 93°C for 3 minutes and slowly cooling down in C1000 Touch™ Thermal Cycler (Bio-Rad). The pre-annealed RNA-TS (32 µM) was mixed with 50 µg of *Pfu*RNA polymerase (4.3 µM) in a 3:1 molar ratio respectively and the mixture was incubated at 80°C for 15 minutes at 80°C. Finally, the Non-Template Strand (NTS) was added to the mixture with a 3:1 molar ration with respect to the polymerase concentration and incubated again at 80°C for 10 minutes. To separate the transcription elongation complex from the excess of nucleic acids, we used a NICK column (GE Healthcare), and fractions were collected manually. The presence of the complex was assessed by 4-15% gradient SDS-PAGE using Mini-PROTEAN® TGX™ Precast Protein gels (BIORAD). Concentration of the RNAP in these fractions was determined by absorbance at 280 nm using NanoDrop™ One/OneC Microvolume UV-Vis Spectrophotometer (Thermofisher) and the most concentrated fraction (0.6-0.7 mg/ml) were used for transcription assays. The transcription reactions were performed as a time courses in the absence of Spt4/5, with Spt4/5 wild-type and the Spt4/5 mutant. The first reaction mixture contained 1 µl of TEC, 3 µl EMSA Buffer 5x (200 mM HEPES/KOH pH 7.4, 300 mM NaCl, 2.5 mM MgCl_2_ x 6 H_2_O, 0.5 mM EDTA, 0.1 mg/ml BSA), and DEPC water 0.1% v/v to arrive to a final volume of 13 µl. 1.5 µl from the nucleotide mix (ATP/CTP 1 mM, UTP 0.02 mM stock solution) was added just before incubation at 80°C. The reaction was stopped through ion chelation by adding 0.5 µl EDTA (0.5 M stock solution) after 30 s, 1 min, 5 min, 10 min, and 30 min. The samples were denatured by addition of 15 µl of 100% formamide containing 0.05% bromophenol blue and incubation at 95°C for 10 minutes. The other two reactions were prepared the same way with the difference that 1 µl of Spt4/5 wild-type or Spt4/5 mutant was added corresponding to a final concentration of 1.7 µM. For each reaction, a control was also prepared which consisted of the same reaction mixture, but the nucleotides were omitted from these samples. The elongation product was evaluated by running an 8 M urea-PAGE (with 20% v/v acrylamide concentration) at 50°C constant temperature and 25 W. The gel plates were 16 x 20 cm and 0.4 mm width and the gel was loaded with 8 µl of the denatured sample. Detection of the Cy5 fluorophore was done using the ChemiDoc™MP Gel Imaging System (BIORAD) and band quantification was also performed using the software ImageLab (BIORAD). To evaluate the Spt4/5 effect on elongation the runoff product from the reaction without Spt4/5 from the 30 minutes sample was equaled as 100% band intensity in each gel. The experiment was performed 4 times.

### Single-molecule FRET studies

The production of double-labelled RNAP from *Methanocaldococcus jannaschii* carrying a fluorescent donor and acceptor dye pair as well as a biotin label in subunit Rpo3 was described in detail elsewhere. The expression and purification protocol to produce recombinant *M. jannaschii* Spt4/5 was reported in detail in the following publications. Briefly, measurements on immobilized RNAP molecules were carried out using a homebuilt PRISM-TIRF setup based on an Olympus IX71 microscope equipped with alternating laser excitation. All measurements were carried out at room temperature (21°C) using a measurement chamber passivated with PEG. Passivated quartz slides and measurements chambers were prepared according to. For excitation, a yellow laser (568 nm; Coherent Sapphire 100 mW) was used for excitation of the donor, and a red laser (639 nm; Toptica iBeam Smart 150 mW) was used for direct excitation of the acceptor, respectively. The fluorescence was collected using a 60x Olympus 1.20 N.A. water-immersion objective. Emission light was split by wavelength using a dichroic mirror (640 DCXR, Chroma Technology) into two detection channels that were further filtered with two bandpass filters (Semrock BrightLine 582/75 and Semrock Brightline 609/54) in the orange channel and one longpass filter (647 nm Semrock RazorEdge) in the near infrared detection range. Both detection channels were recorded by one EMCCD camera (AndorIX on X3, preGain 5.1, gain 250, frame rate 10 Hz) in a dual view configuration (TripleSplit, Cairn UK) and the acquired data were analyzed by custom-made software based on LabVIEW 2012 64bit (National Instruments). Data analysis was furthermore carried out using the software Origin. The resulting histograms were fitted either with a single or double Gaussian fit and the mean FRET efficiency and the standard error was determined from the fit. Spt4/5-RNAP complexes were pre-formed at 65°C for 20 minutes prior to immobilization on the streptavidin-coated surface and subsequent microscopy. Similarly, RNAP apo enzyme was incubated for 20 minutes at 65°C without addition of Spt4/5 prior to single-molecule measurements.

## Results

### Cryo-EM reconstructions of apo RNA polymerase from *P. furiosus* show the clamp/stalk module in expanded and contracted states

In this work, we purified the native archaeal RNAP from the hyperthermophilic euryarchaeon (Methanobacteriota) *Pyrococcus furiosus* (**Supplementary Figure S1**). To this end, a mutant *Pfu* strain constitutively expressing C-terminally Strep-and His-tagged RNAP subunit Rpo3 from its native genomic locus was created, allowing affinity purification of the cellular archaeal RNA polymerase (**Figure S1**).*Pfu* RNAP is used as a standard model system for archaeal transcription and shows high sequence identity (83.8%)with the archaeal RNAP from *Thermococcus kodakarensis*, of which structures were analyzed before^45,48^.Cryo-EM data was collected on a CryoARM200 (JEOL) equipped with a GatanK2 direct electron detector and in-line energy filter and processed using the RELION suite. Single-particle reconstructions of the apo form of the RNAP from *Pfu* RNAP were determined at final resolutions of up to 3.2 Å (**Figure 1**, **Figure S3, Figure S4, Table S1**). Damaged, oligomerized and undefined particles were removed during data processing (**Supplementary Figure S1**). *Pfu* RNAP is composed of 11 subunits devoted to catalysis (Rpo1N, Rpo1C, and Rpo2), assembly of the RNAP (Rpo3, Rpo10, Rpo11, and Rpo12), and auxiliary functions (Rpo4, Rpo7, Rpo5 and Rpo6) (**Supplementary Figure S2**). The reconstructions reveal well-defined cryo-EM density for all 11 *Pfu*RNAP subunits, allowing model building starting from *Thermococcus kodakarensis* RNAP models^45,48^. A reconstruction containing 161,531 particles shows the RNAP in an open-clamp conformation characterized by an expanded central DNA-binding cleft spanning ∼54.3Å from the clamp core helices to the protrusion domain (**Supplementary Figure S6a**). The DNA-binding cleft is not occupied by any density, indicating that complete purification of residual nucleic acids was successful and an inactive apo-RNAP reconstruction was obtained. Central functional elements such as the bridge helix and the catalytic Mg^2+^ ion are well ordered and show defined density, while residue positions in the trigger loop remain flexible (**Figure 1a,d)**. However, local resolution estimation in RELION indicated dynamic conformations of the stalk, clamp head and clamp core, as well as the jaw domain within subunit Rpo1C.To analyze the structural heterogeneity of these domains, we employed the variability analysis included in CryoSPARC^52^, confirming the dynamics of the clamp/stalk (**Supplementary Movie 1**) and jaw regions (**Supplementary Movie 3**). Furthermore, this allowed the reconstruction of an additional contracted conformation at a resolution of 3.0 Å (**Figure1b** and **Supplementary Figure S4**). These results indicate that the *Pfu*RNAP exists in a continuum of structural conformations within an expanded and a contracted boundary in its apo form *in vitro*.

This is well in line with the results of single-molecule FRET studies using the recombinant archaeal RNAP from the related archaeal organism *Methanocaldococcus jannaschii* showing that the clamp domain can adopt an open and closed conformation in the apo state RNAP, with additional molecules adopting a conformation in between the closed and open clamp state suggesting that the clamp is in a dynamic equilibrium between the open and closed state.

### Spt4/5 binds the RNA polymerase in the absence of nucleic acids and induces a conformational change

In the same cryo-EM dataset of the apo *Pfu* RNAP, about half of well-defined particles showed additional cryo-EM density spanning the DNA-binding cleft between the clamp core helices and the protrusion domain. Following 3Dclassification and refinement in RELION, a 3D reconstruction of these particles at an overall resolution of 3.4 Å was obtained (**Figure 1c, Supplementary Figure S3** and **Table S1**). Using the model of a recombinant clamp-Spt4/5 crystal structure as reference, we were able to unambiguously attribute the additional density to Spt4/5^21^. The position overlaps with interpretations of low-resolution EM-envelops. Density for the Spt5 KOW domain was not observed in our reconstructions, indicating that the domain is flexibly linked to the NGN in this state and the KOW domain has rotational freedom. Local resolution differences (**Supplementary Figure S4**) and flexibility analysis in cryoSPARC indicate that the position of Spt4/5 is not rigidly defined (**Supplementary Movie 4**). The RNAP appears to be in a ‘super-contracted’ conformation when bound by Spt4/5, meaning that the clamp domain closes beyond the most contracted apo RNAP conformation we observed (**Supplementary Movie 4**). In the super-contracted conformation, the stalk sub-complex and the jaw domain contract alongside the clamp domain. Furthermore, theSpt5-NGN domain reaches so far across the DNA-binding cleft that it contacts the lobe in addition to the protrusion domain.

Note that, strikingly, the DNA binding channel of the RNAP is not accessible for DNA and that initiation factor binding sites are occluded in the apo-Spt4/5 configuration, preventing productive initiation or elongation (**Supplementary Movie 2)** and Spt4/5 would have to dissociate to allow loading of the DNA during the initiation phase of transcription.

To verify that endogenous Spt4/5 indeed co-purifies with *Pfu* RNAP from cellular extracts, we performed mass spectrometry (MS) analysis of the affinity purified RNAP (**Supplementary Figure S5**). In addition to all RNAP subunits, Spt5 was also enriched in these samples (**Figure 2a**). Spt4 was slightly less enriched, likely owing to the small size of the protein. In addition to MS analysis, the co-purification of Spt4/5 with RNAP was confirmed by Western Blot analysis using an antibody directed against *Pfu* Spt4/5 (**Figure 2b**).

**Figure 2:**
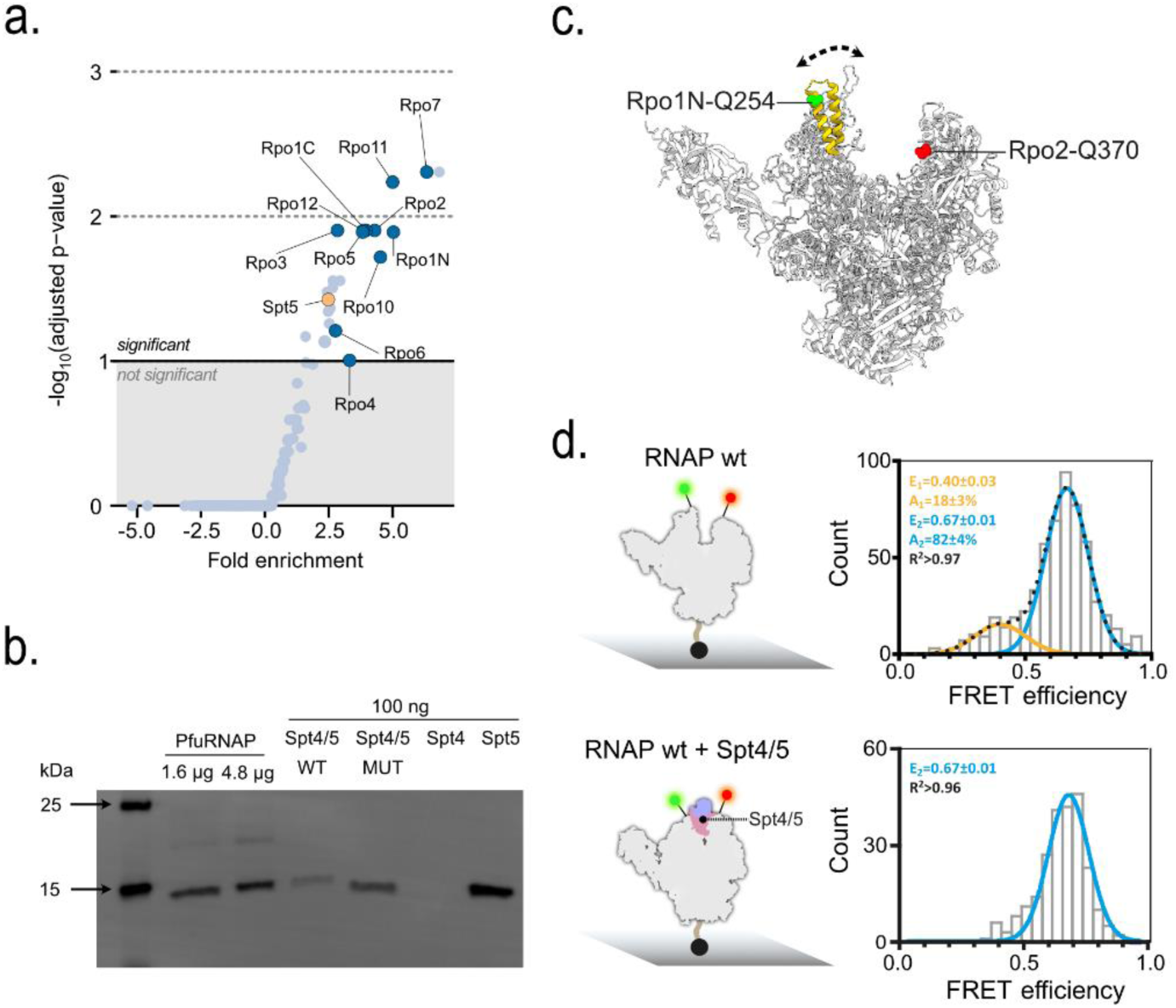
Accessing the RNAP-Spt4/5 interaction using mass spectrometry and single-molecule FRET measurements. **(a)** Volcano plot of mass spectrometry analysis conducted for the *Pfu* rpoD mutant with respect to the negative control (“His-tag only” mutant). Fold enrichment over background was calculated using a one-sided Wald test with multiple testing correction using Benjamin-Hochberg. RNAP subunits and Spt5 are color-coded in blue and yellow, respectively. **(b)** Western blot analysis of samples derived from affinity-purified *Pfu*RNAP from a cell lysate from a *Pfu* variant that expresses His-tagged subunit Rpo3. Recombinantly expressed and purified wild-type or mutant Spt4/5 (100 ng) as well as the individual recombinant proteins Spt4 and Spt5 (100 ng) were loaded as reference. For detection, an antibody raised against the elongation factor Spt4/5 was used. **(c)** Structure of the 12-subunit archaeal RNA polymerase from *Pyrococcus furiosus*(PDB: 8ORQ) with an open clamp (yellow). The archaeal RNA polymerase from *Methanocaldococcus jannaschii* was site-specific labelled with a donor (DyLight550) and acceptor fluorophoreDyLight650 in subunit Rpo1N*^E257^(green)and the protrusion domain (subunit Rpo2C*^Q373^, red), respectively. For immobilization purposes a biotin was coupled to a native cysteine in subunit Rpo11 (black sphere protruding from the RNAP core). (**d**) FRET efficiency histograms show smFRET efficiencies determined for the wildtype RNA polymerase (RNAP wt) and in complex with transcription factor Spt4/5 (RNAP wt + Spt4/5). All measurements were performed in the absence of nucleic acids. The histograms were fitted with a single or double Gaussian function and the mean FRET efficiencies (E), the percentage distribution of the populations (A) and the coefficient of determination (R^2^) are given with standard errors in the histograms.

We next asked whether the different conformational states of the RNAP and RNAP in complex with Spt4/5 can also be monitored in solution. Complementing the cryo-EM analysis, we made use of our previously established single-molecule FRET assay using the RNAP of the closely related archaeon *M. jannaschii*. This assay reports on the distance and the distance changes between a fluorescent donor-acceptor dye pair engineered into two residues that are located opposite of each other across the DNA-binding cleft (**Figure 2c**, residue 257 in Rpo1Nand residue 373 in Rpo2C, respectively - numbering for the RNAP of *M. jannaschii*).The *M. jannaschii* RNAP is found in an open (low FRET with mean FRET efficiency of E = 0.40) and closed clamp state (low FRET with mean FRET efficiency of E = 0.67) in solution (**Figure 2d**) reflecting the conformational states found in our cryo-EM studies. Our data furthermore reveal that addition of Spt4/5 shifts the equilibrium towards the population with high FRET efficiency(**Figure 2d**)Even though we observed a further closure of the clamp in the supercontracted RNAP state, we do not observe a shift in mean FRET efficiency, which is the same as for the unliganded RNAP (E = 0.67).However, a distance change of 2 Å between contracted and supercontracted state would only result in a marginal change in FRET efficiency of 0.03, which is too small to be reliably detected. Taken together, these data further support the finding that binding of Spt4/5 to the apo RNAP is possible even in the absence of DNA and that Spt4/5 induces conformational changes in RNAP.

### The structure of an archaeal transcription elongation complex is similar to eukaryotic counterparts, with some exceptions

To investigate the three-dimensional architecture of the archaeal TEC, *Pfu*RNAP was recruited to a synthetic elongation scaffold following protocols described for TECs formed with the RNAP from *Methanocaldococcus jannaschii*^45,48^ and subjected to single-particle cryo-EM analysis (Methods, **Figure 3a, Supplementary Figure S8, Table S1**). Following processing and 3D reconstruction in RELION, we obtained a Cryo-EM map of the TEC at an overall resolution of 3.0 Å. In addition to the RNAP subunits, the density allowed us to build a molecular model for the downstream DNA double helix and the DNA-RNA hybrid, enabling investigation of nucleic acid interactions within the TEC (**Figure 3 c/e**). The model includes nine ribonucleotides of the RNA, but local resolution at the areas of the RNA-exit channel and within the center of the bubble prevented further RNA and non-template strand (NTS) modelling. Comparable to other RNAP-TEC^8,53^ reconstructions and crystal structures, the downstream dsDNA double helix was well-defined, whereas the upstream dsDNA region showed higher flexibility. The primary contacts between the RNAP and nucleic acids are not sequence-specific but involve several basic residues and the DNA phosphate backbone (**Figure 3c**). The RNAP is found in the pre-translocated state as indicated by the catalytic magnesium contacting the +1 ribonucleotide. The bridge helix separates position -1of the template strand from -2 on the DNA template strand and is found in a fully helical “unstretched” state. Overall, the structure of the elongation complex shows high similarity to the three-dimensional organization of the eukaryotic TEC containing RNAP II^54^ (**Supplementary Figure S11**). This includes the clamp/stalk module that adopts the contracted state, securing the downstream DNA in the DNA binding channel (**Figure 3**, **Supplementary Movie 1)**. However, a conserved contact of the downstream DNA duplex with the subunit Rpb5,as observed for eukaryotic TECs, is not reflected in the archaeal TEC because the N-terminal domain of eukaryotic Rpb5 is not conserved in Archaea. Structural comparison of the *Pfu* TEC reconstruction with the Spt4/5-engaged apo polymerase showed many common features, such as the overall contracted RNAP state with closed clamp domain, contracted stalk sub-complex and jaw domain. This leads to the conclusion that the RNAP conformation would remain identical when Spt4/5 subsequently engages with the TEC. To test this hypothesis, we again employed single-particle cryo-EM to determine a reconstruction of the Spt4/5-decorated TEC.

**Figure 3:**
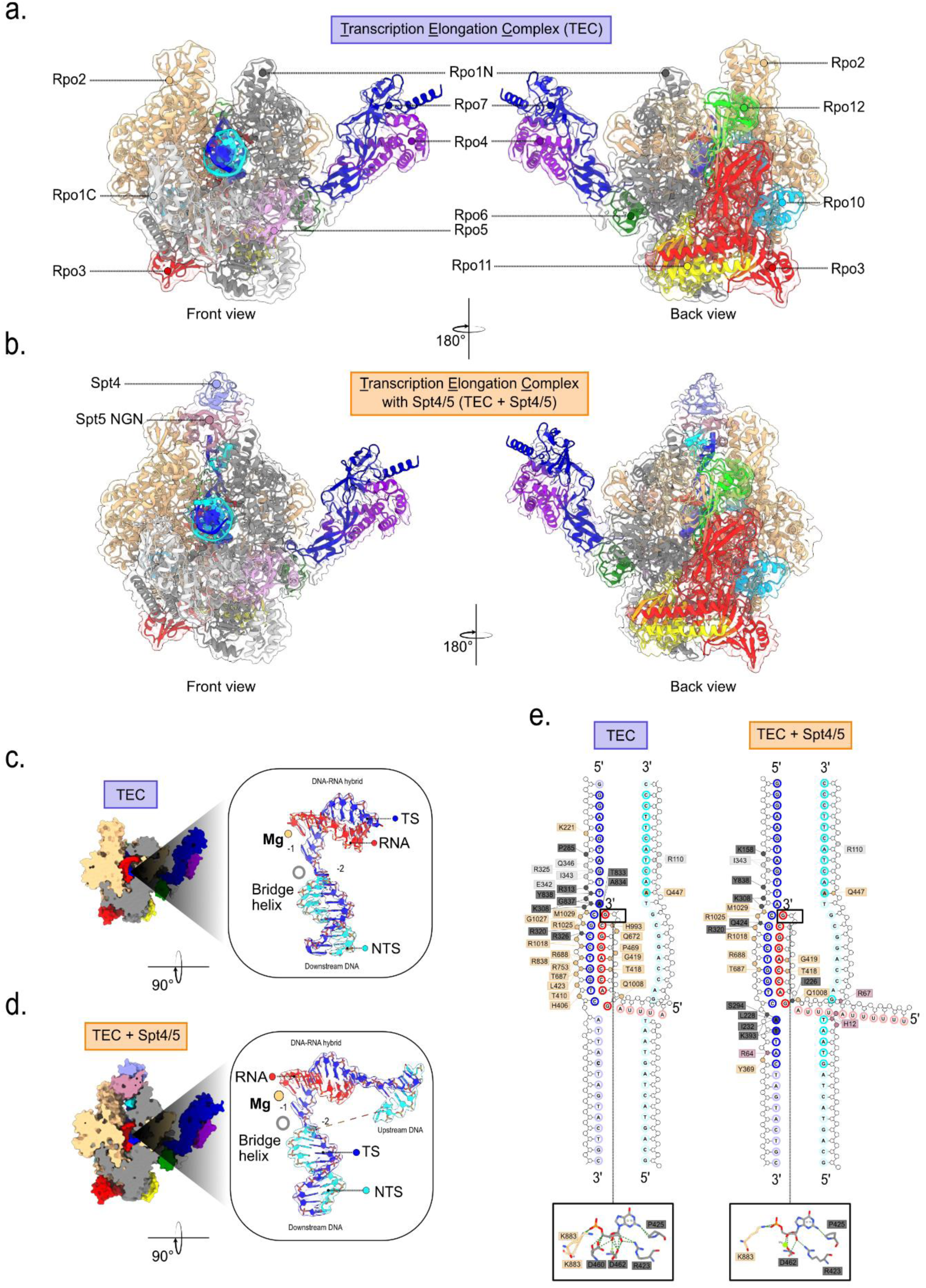
Structure of the euryarchaeal transcription elongation complex in presence and absence of Spt4/5. Structural model of the TEC in **(a)** the absence and **(b)** presence of Spt4/5 in two views. **(c)** Schematic representation of the nucleic acid scaffolds used for the TECs formed in the absence (top) and presence (bottom) of Spt4/5. Template strand colored in blue, non-template strand in cyan, RNA in red. Interactions with of amino acids side chains with the nucleic acid scaffold are indicated. Coloring of the RNA polymerase subunits as in (a). **(d)** Nucleic acid strand model in the TEC and the TEC in the presence of Spt4/5. The magnesium cation in the metal A position is shown as orange ball. The position of the bridge helix is indicated with a grey sphere. Part of the downstream DNA can be traced when Spt4/5 is part of the TEC. **(e)**Schematic representation of *Pfu* RNAP and nucleic acids contacts calculated in ChimeraX, in the TEC in absence (left) and in presence (right) of Spt4/5.

Notably, the C-terminal region of Rpo2 (residues 1005 to 1024) that contacts the DNA-RNA hybrid appears to be fully flexible within the open RNAP form but adapts a fully folded and active conformation within the contracted form in a conformation that is identical to the TEC and Spt4/5-containing TEC. This stretch binds to the DNA-RNA hybrid, specifically the phosphate backbone of the template strand in both TEC reconstructions.

### Archaeal Spt4/5 engages transcribing and non-transcribing RNAP in different orientations

To determine the structure of the archaeal TEC in complex with Spt4/5, the complex was formed by adding recombinantSpt4/5 to a pre-formed TEC in excess (methods) followed by stabilization using chemically crosslinking with BS^3^ prior to SEC to remove incompletely assembled particles (**Supplementary Figure S9 and S10**).Here, we used an elongation scaffold containing a longer RNA since structural studies of RNAP II in complex with Spt4/5^55,56^ showed contacts emerging RNA via its KOW domains. Again, we collected cryo-EM data (methods, **Supplementary Figure S9, S10**). Following processing and single-particle analysis, a reconstruction at an overall resolution of 3.4Å resolution was obtained, allowing to construct a molecular model of the Spt4/5-containing TEC (**Figure 4b** and **Supplementary Figure S10**, **Table S1**).Even though the local resolution is reduced in the region where Spt4/5-interacts with the RNAP clamp, an adaptation of the previous model showing Spt4/5 in contact with contracted apo RNAP (see above) allowed the construction of a confident model. Nevertheless, consistent density for the KOW domain was absent, indicating that the domain is either flexibly linked to the NGN domain or subject to positional heterogeneity in the archaeal TEC. Contacts between the Spt5 NGN domain and the RNAP clamp and protrusion domains are established, as shown in **Supplementary Figure S11a**. In comparison to the previous TEC reconstruction, the Spt4/5-containing TEC shows clear density for the upstream dsDNA duplex, allowing to construct a model for this part and hinting at a stabilization of this DNA part by Spt4/5 within the TEC (**Figure 3d**). Furthermore, comparison of the position of Spt4/5 within the TEC and in the apo RNAP reveals that helix 1 of Spt5 is found in a position shifted by ∼6 Å (**Supplementary Figure S11**). In context of the TEC, Spt5 helix 1 is shifted away from the protrusion domain and thus partially retracts from the DNA-binding cleft of the RNAP. This repositioning is likely required to accommodate DNA within the central cleft. Upon partial retraction in the TEC, contacts of Spt5 with the RNAP lobe domain cannot be established any longer, preventing the transition into a super-contracted state as found in the nucleic acid-free RNAP complexed with Spt4/5.

**Figure 4:**
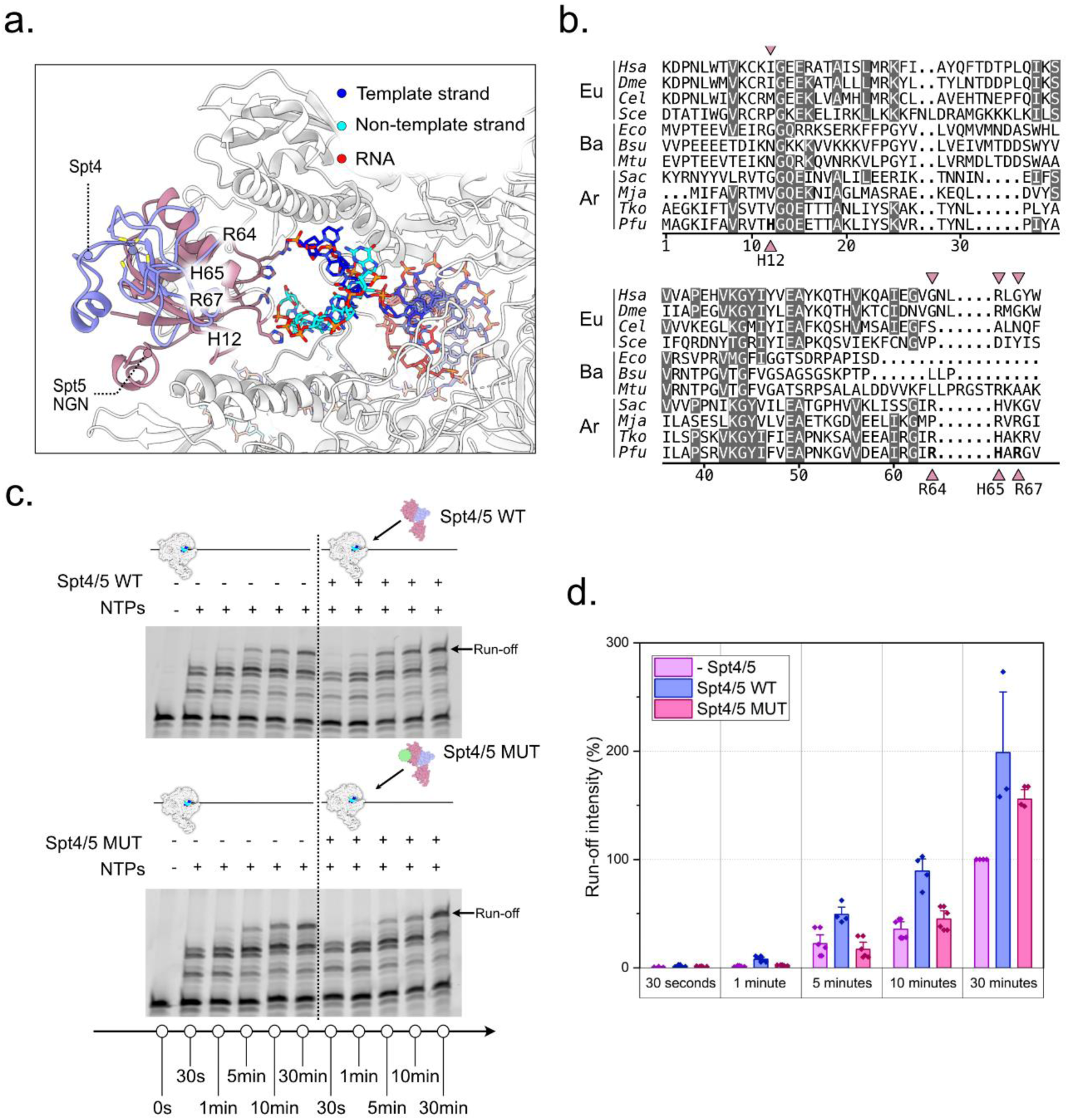
Interactions between Spt4/5 and the non-template strand within the upstream DNA duplex enhance the transcriptional activity of the archaeal RNA polymerase. **(a)** Close-up view of the interaction of Spt5 with the phosphate backbone of the non-template strand. Spt5 sidechains (H12, R64, H65 and R67) that contact the DNA are indicated. Spt5 is colored in dusky pink and Spt4 in purple. **(b)** Multiple sequence alignment of Spt5 including eukaryotic Spt5 variants (*Hsa*: *Homo sapiens*, *Dme*: *Drosophila melanogaster*, *Cel*: *Caenorhabditis elegans*, *Sce*: *Saccharomyces cerevisae*), bacterial NusG variants (*Eco*: *Escherichia coli*, *Bsu*: *Bacillus subtilis*, *Mtu*: *Mycobacterium tuberculosis)* and archaeal Spt5 variants (*Sac*: *Sulfolobus acidocaldarius*, *Mja*: *Methanocaldococcus jannaschii*, *Tko*: *Thermococcus kodakarensis*, *Pfu*: *Pyrococcus furiosus*). Conserved residues are highlighted in grey. The residues that interact with the non-template strand in the TEC of *Pfu* are indicated with a triangle. **(c)** Transcription elongation assay using wt *Pfu*RNAP (0.6-0.7 µg) and Spt4/5 wt or the Spt4/5 mutant (Spt5^H12A/R64A/H65A/R67A^) (1.7 µM). As references, transcription assays in the absence of Spt4/5 were performed. Reactions were stopped at 0.5, 1, 5, 10 and 30 min. (d) Diagram showing the quantified run-off transcript levels shown in panel (c). Bar plots represent normalized RNA band intensities (mean +/- SD) from three technical replicates.

### Spt5 apparently stabilizes the upstream DNA duplex

Contacts between the NGN domain ofSpt5 and the backbone of the upstream DNA duplex in the TEC reconstruction were putatively identified, involving the four residues His12, Arg64, His66, and Arg67 (**Figure 4a**). Although the precise sidechain orientations in this contact region remain partially defined, our analysis indicates that the presence of Spt4/5, compared to the free TEC, leads to the fixation of the transcription bubble at its upstream edge, stabilizing the upstream duplex.

To assess the significance of DNA engagement by the four identified basic residues for Spt5 function, we conducted a standardized *in vitro* transcription assay. The results demonstrate that the presence of wild-type Spt4/5 prevents pausing and stimulates the processivity of *Pfu*RNAP (**Figure 4c/d and Supplementary Figure S13**).However, the stimulatory effect of Spt5 was reduced in an HRHR-AAAA mutant, generated by mutating the contributing residues in the Spt5-DNA interface, even though it was still able to stably bind Spt4 as concluded from heterodimer formation during the purification process (**Figure 4a**).

Interestingly, the four basic Spt5 residues that contact the upstream DNA are not conserved across archaeal phyla or when compared to bacterial NusG or to eukaryotic Spt5 (**Figure 4b**). Taken together, the results of this structure-function analysis indicate that the interaction between Spt5 and the upstream edge of the transcription bubble is essential for Spt5 function in *Pfu*.

## Discussion

### Evolutionary aspects regarding clamp movement and cleft contraction

The clamp domain is a mobile element crucial to the functioning of all multi-subunit RNAPs, as shown by a wealth of structure-function and single-molecule FRET analyzes. However, there is no consistent picture regarding the conformation of archaeal RNAPs in their apo form, as the clamp was found in either closed or open states. In this study, we fill gaps in the understanding of the conformational status of the clamp in Archaea, bringing together previous observations of archaeal RNAP clamp flexibility in the apo state and clamp fixation upon transition to active elongation. Single-particle cryo-EM reconstructions of *Pfu* RNAP presented in this work showed the enzyme in expanded and contracted conformations that are defined by different positions of the clamp domain and the stalk sub-complex (**Figure 1a,b**). Three-dimensional variability analysis revealed a high degree of clamp/stalk module flexibility among all particles, which is well in line with single-molecule FRET (smFRET) measurements showing that expanded and contracted conformations can both be adopted in solution, suggesting that the clamp is fully flexible when the RNAP is not engaged with the template DNA or transcription factors. In the elongation state, the clamp closes, and the DNA-binding cleft is found in a contracted state (**Figure 2a,b**).Well in line with our findings, another parallel work elucidated the structural organization of an archaeal pre-termination complex, showing the clamp in a contracted conformation before termination takes place^57^. It is assumed that the clamp reopens to release the DNA and RNA during termination^39,58,59^.

Similarly, functional clamp movements have been observed for bacterial and eukaryotic RNAPs^6,7^. Structural and single-molecule analysis of bacterial RNAPs at different steps of the transcription cycle elucidated the mobile nature of the clamp domain. For example, the bacterial apo RNAP exhibits an open clamp in crystal structures and cryo-electron microscopy structures^60,61^. Time-resolved single-molecule FRET measurements of the RNAP apo enzymes from *E. coli* moreover revealed that the clamp is conformationally flexible and populates three distinct states^62^. In bacterial transcription elongation complexes^35,63^, the clamp adopts a closed state and apparently reopens during transcription termination^64^. These clamp movements are of functional importance for bacterial but also archaeal transcription as an opening of the clamp promotes for example transcription initiation^19,65^ most likely by supporting the separation of the DNA strands. Clamp flexibility was not observed for apo RNAPs from the archaeal Thermoproteota phylum. For this phylum, crystal structures (*S. shibatae* and *S. solfataricus*)^47,50^ and a cryo-EM structure from *S. acidocaldarius*^66^ were reported and in all cases a closed clamp conformation was found for the RNAP apo form. The viral RNAP-interacting protein ORF145 and TFS4 were shown to impact the *S. acidocaldarius* RNAP clamp conformation and to shift the clamp to a more open state as compared to the apo enzyme (**Supplementary Figure S6b,e**). Nevertheless, the crystal structure^45^ of *T. kodakarensis* RNAP was resolved in an expanded conformation and in a contracted conformation in the cryo-EM structure^48^, similarly to *Pfu* RNAP apo forms. FRET and structural studies from *P. furiosus,T. kodakarensis* and *M. jannaschii* suggest a mechanistic conservation of a flexible clamp of the apo RNAP throughout the archaeal Methanobacteriota phylum members, which could not be verified in RNAPs from the Thermoproteota phylum to date.

Opening and closing of the clamp domain was also observed in eukaryotic RNA polymerases^39^, including RNAP II of yeast and mammalian origin. However, clamp movements were documented to be more pronounced for mammalian RNAP II. For example, density for the clamp was found to be weak or absent in a cryo-EM datasets for the RNAP II associated with protein 2 (RPAP2), which binds between the jaw domains of RNAP II^54,67^ in absence of nucleic acids. Similarly, the clamp is highly mobile in the dimeric RNAP II structure from *S. scrofa domesticus*^68^. Structures of the transcribing bovine RNAP II showed that even in the TEC a small subfraction of the RNAPs (10%) exhibit a mobile clamp in the absence of Spt4/5. Notably, the activity of RNAP I relies on modulation of its overall contraction state, including the clamp in inactive conformations, a full expansion of the clamp-stalk module occurs, even allowing RNAP dimerization^6,7^. The RNAP is found in a fully contracted state upon initial transcription^69^ and remains contracted throughout productive elongation, as shown by single-particle cryo-EM analysis and tomography analysis^70–72^. The apo form of RNAP I may also adapt different clamp contraction states. Especially in human RNAP I, clamp flexibility is strongly pronounced, indicating a conformational continuum^73^. Hence, *Pfu*RNAP and eukaryotic RNAP I appear to undergo similar expansion/contraction movements.

### Spt4/5-engagement limits conformational flexibility of the archaeal RNAP

Interaction of bacterial NusG with DNA-free bacterial RNAP has been reported^74^, but a comparable interaction between Spt4/5 and the DNA-free state of RNAP was never observed for eukaryotic RNA polymerases. This might be because a tight interaction between eukaryotic Spt4/5 and the RNAP requires the presence of the newly synthesized RNA that forms an interaction with the linker domain between KOW_4_ and KOW_5_ domain of eukaryotic Spt5^55,56^. Here, we conclude that archaeal Spt4/5 binding does not strictly require KOW-interactions with the product RNA and is only possible when the RNAP adopts the closed clamp conformation (**Figure 2b**). Likely, and well in line with smFRET analyses *in vitro*, archaeal RNAP adapts various states in a dynamic equilibrium within the contracted and expanded boundaries we describe. Spt4/5 apparently selects for a specific conformational state and can only bind contracted RNAPs, resulting in super-contraction and removal from the conformational flexible pool of apo RNAPs. However, we cannot rule out that Spt4/5 binding induces the closure of the clamp instead. For Spt5 from *M. jannaschii*, residues A4 and Y42 are of crucial importance for the interaction with the RNAP^20^. These residues are conserved in *Pfu* and similarly contribute to the interaction with the RNAP clamp via its hydrophobic pocket together with a hydrogen bond between Spt5 E78 and Rpo1N Q262, previously observed in the crystal structure^21^.

Loading of the DNA into the binding channel requires an open state of the clamp, a structural transition supported by TFE during transcription initiation^19,48^. Our cryo-EM data suggest that a subfraction of the cellular RNAP pool is associated with Spt4/5 in the absence of a nucleic acid substrate. In the *Pfu* RNAP-Spt4/5 complex, the clamp is closed, and the DNA binding channel blocked by Spt4/5. Consequently, DNA cannot enter the RNAP in an Spt4/5 pre-engaged state. This may indicate a mechanism to prevent binding of unspecific DNA. Similar strategies are employed in RNAP I and III using intrinsic structural elements (specifically, the C-terminus of subunit RPC7 in RNAP III^75^ or the DNA-loop within the largest subunit of RNAP I^6,7^). RNAPs decorated with Spt4/5 might represent an RNAP mimicking ‘expander’ reservoir available in the cell that can quickly be activated. To activate *Pfu* RNAP molecules for promoter-directed transcription, Spt4/5 must be displaced from the RNAP. Spt4/5 is known to compete with TFE for the clamp^11,17^ leading to a replacement of TFE during the transition from initiation to elongation. A reverse scenario, e.g. the displacement of Spt4/5 by TFE to prime the RNAP for transcription initiation, might allow the activation of this RNAP pool under circumstances that demand higher transcriptional output or a switch of the transcriptional program. In line with this idea, we have previously shown that TFE has a higher affinity for the RNAP in context of the PIC as Spt4/5^17^. A somewhat similar mechanism is employed by the viral protein ORF145 which occupies the DNA binding channel of the RNAP and effectively inhibits productive transcription^49^. As the respective virus does not encode its own RNAP, release of ORF145 from the archaeal RNAP must take place to allow the transcription of the viral genome. The mechanism by which this is achieved, however, is unknown. Compared to the RNAP-ORF145 structure, the extent of RNAP clamp contraction is more pronounced in the Spt4/5-TEC (**Supplementary Figure S6**). For these reasons, we hypothesize that the reversible shutdown of RNAP activity by temporary fixation of the cleft contraction state may be a regulatory strategy.

### The Spt5-interface with upstream DNA in TECs is functionally important in *P. furiosus*

Bacterial NusG and archaeal Spt5 (in complex with Spt4) are well known for their function as transcription elongation factors that stimulate the processivity of the RNAP^4^. The underlying mechanism for this stimulatory activity is found in the overall architecture of the TEC with Spt4/5 and NusG, which bridge across the DNA binding channel to secure the DNA. Additionally, Spt5/NusG prevents structural changes in the RNAP that could otherwise lead to backtracking or termination. Specifically, it is suggested that Spt4/5 ensures that the clamp remains in a firmly closed state. In eukaryotic complexes, a contact between RNA and Spt5 is formed by the linker domain between KOW_4_and KOW_5_ domain, which is not present in archaeal Spt5^22^. Our reconstructions show that the overall architecture and organization of the TECs is conserved between Bacteria, Archaea and Eukaryotes, even though DNA contacts may vary among organisms and transcription systems. We do, however, not observe direct contacts of the Spt5 KOW domain with the product RNA, even though this may depend on the scaffold and could change in a state in which the Spt5 KOW domain adopts a defined position.

Our structures are also well in line with results of single-molecule FRET studies that show that the conformational equilibrium between a closed and intermediate state is shifted towards the intermediate conformation in the presence of Spt4/5.

Interestingly, while NusG has anti-pausing activity in *E. coli* like archaeal Spt4/5, no contacts between NusG and the NTS were reported so far for the TEC in *E. coli*. A multiple sequence alignment showed that the residues H12, R64, H65 and R67 are not highly conserved among NusG and Spt5 homologous. Notably, the histidine at position 12 is not even conserved among archaeal species and appears to be a unique residue in the NGN domain of Spt5 from *Pfu*. Mutating these residues led to a reduction of Spt4/5-dependent stimulation of RNAP’s processivity (**Figure 4c,d**). We conclude that the Spt5 residues in the upstream-duplex interface are important for the function of Spt5 in *Pfu,* suggesting that archaeal Spt4/5 stimulates the processivity of the RNAP in a dual manner: i) by its role as bridging factor that locks the DNA in the binding channel and ii) by an interaction of the NGN domain with the upstream duplex to stabilize the transcription bubble. Similarly, the transcription initiation factor TFE stabilizes the upstream edge of the transcription bubble in initially transcribing complexes, demonstrating that this is crucial to the integrity of transcription complexes^76^.Most likely, such a stabilization by Spt5 of TFE is of even greater importance in hyperthermophilic Archaea as DNA melting and thus RNAP-dissociation is favored by the high growth temperatures under which these organisms thrive. In this respect, TFE and Spt5 do not only physically but also functionally replace each other upon the factor-swapping mechanism in the transition from initiation to elongation^77^. To which extent the interaction between the NTS and Spt5 is conserved across Archaea, however, remains to be determined by solving for example the elongation complex structures of well-known archaeal model systems like *Sulfolobus* or *Saccharolobus* species or from *Thermococcus kodakarensis*. For *T. kodakarensis* it was shown that Spt4/5 is crucial for the recruitment of the archaeal termination factor aCPSF1/FttA^37,38^. Spt4/5 directly interacts with one of the aCPSF1 copies via residues F106, V131, V132 and I134. Moreover, aCPSF1 forms contacts with the stalk domain and the RNA. The KOW domain is not resolved in our structure indicating that this is a highly flexible domain that is stabilized by the interaction with aCPSF1. While only residues V131 is conserved between Spt5 from *Pfu* and *T. kodakarensis*, the overall fold of Spt5 homologous is highly similar, indicating that similar interactions can be formed in *Pfu*.

Taken together, we provide structural insights into archaeal transcription, which highlight the high degree of conformational flexibility of the clamp domain of the archaeal apo RNAP enzyme. We reveal that the apo RNAP-Spt4/5 complex adopts a super-contracted state, which hints to the existence of an inactivated RNAP pool in the cell. We furthermore provide the structure of an archaeal transcription elongation complex in the absence and presence of Spt4/5, revealing interactions of the Spt5-NGN domain with the upstream DNA, which are relevant for the function of Spt4/5 as an elongation factor.

## Data availability

The datasets generated during and/or analyzed during the current study are available from the corresponding author on reasonable request. Structural data are deposited at the protein data bank: *Pfu* expanded apo RNA polymerase 8ORQ, EMD-17130; *Pfu* contracted RNA polymerase 8RBO, EMD-19033; *Pfu* RNA polymerase in complex with Spt4/5 8P2I, EMD-17366; Transcription Elongation complex 8CRO, EMD-16809; TEC bound to Spt4/5 8OKI, EMD-16929. The mass spectrometry proteomics data have been deposited to the ProteomeXchange Consortium via the PRIDE partner repository with the dataset identifier PXD048268

## Acknowledgements

We would like to thank Sarah Schulz for single-molecule FRET measurements and analysis and Finn Werner for discussions regarding smFRET experiments. We would furthermore like to thank Prof. Dr. Ralph Witzgall and Prof. Dr Reinhard Rachel for JEM2100F access and training. Some cryo-EM data were collected at the cryo-EM facility of the University of Würzburg with support from Bettina Böttcher and Christian Kraft. We thank Henning Plikat for support with sample preparation for mass spectrometry analysis. Single Particle cryo-EM – including sample preparation course 2021, organized by Rebecca Thompson. We thank Philipp Becker and Dr. Simona Pilotto for advice in model building, David Pöllmann for the construction of the *Pyrcococus* strain used for RNAP purification, Martin Brehm for protein purification and crosslinking assistance. We thank Dipl. Ing. (FH) Thomas Hader and Simon Dechant for technical assistance to cultivate *Pfu* in large-scale bio-fermenters. Furthermore, we would like to thank all members of the transcription team at the Institute of Microbiology for fruitful discussions. The authors thank all past and present members of the structural biochemistry group within the RCB for their contribution, Norbert Eichner and Gerhard Lehmann for their invaluable IT support and Katharina Vogl and Mona Höcherl for technical assistance.

## Author contributions

**Christoph Engel** designed and supervised (cryo-)EM work, processing and data analysis, model-building and interpretation; contribution to funding, manuscript writing, discussion and editing, figure and movie generation

**Dina Grohmann:** designed and supervised the research, biochemical data analysis, interpretation of *Pfu* RNAP structures, smFRET data analysis and interpretation, expression and purification of labelled RNAP and unlabelled Spt4/5 (*M. jannaschii* transcription system), securing funding, wrote the manuscript and created figures

**Felix Grünberger:** cultivation of *P. furiosus* and initial mass spectrometry analysis of *Pfu* RNAP purified from cell mass, bioinformatical MS data analysis, bioinformatical analysis (multiple sequence alignment) and contribution to figures and material and methods section, discussion of the manuscript

**Winfried Hausner:** creation of *P. furiosus* strain expressing tagged *Pfu* RNAP, *Pfu* RNAP preparation, initial experimental design and experiments for MS analysis of RNAPs, discussion of the manuscript

**Florian Heiß** cloning of *Pfu* Spt4/5 expression plasmids, support in (cryo-)EM sample preparation and processing

**Michael Pilsl** conducted and instructed (cryo-)EM work, sample preparation and optimization, processing and data analysis, model-building and interpretation; contribution to manuscript writing, discussion and editing, figure and movie generation

**Robert Reichelt:** large scale and small-scale cultivation of *P. furiosus* wild-type and mutant strains, support in *Pfu* RNAP and Spt4/5 purification, support in transcription assay development, discussion of the manuscript

**Daniela Tarau:** RNAP purification, RNAP negative staining, sample preparation for cryo-EM, establishing X-linking for TEC, cryo-EM data processing, model building, analysis and interpretation of *Pfu* RNAP structures, purification of Spt4/5 wt, cloning and purification of Spt4/5 mutants, transcription assays, preparation of *P. furiosus* for mass spectrometry analysis, Western Blot analysis, manuscript writing, creation of figures and movies

**Sabine König, Henning Urlaub:** LC-MS analysis

## References

1. Osman, S. & Cramer, P. Structural Biology of RNA Polymerase II Transcription: 20 Years On. Annu. Rev. Cell Dev. Biol. 36, 1–34 (2020).

2. Griesenbeck, J., Tschochner, H. & Grohmann, D. Structure and Function of RNA Polymerases and the Transcription Machineries. in Macromolecular Protein Complexes (eds. Harris, J. R. & Marles-Wright, J.) vol. 83 225–270 (Springer International Publishing, 2017).

3. Werner, F. & Grohmann, D. Evolution of multisubunit RNA polymerases in the three domains of life. Nat Rev Microbiol 9, 85–98 (2011).

4. Gehring, A. M., Walker, J. E. & Santangelo, T. J. Transcription Regulation in Archaea. J Bacteriol 198, 1906–1917 (2016).

5. Chen, J., Boyaci, H. & Campbell, E. A. Diverse and unified mechanisms of transcription initiation in bacteria. Nat Rev Microbiol 19, 95–109 (2021).

6. Fernández-Tornero, C. et al. Crystal structure of the 14-subunit RNA polymerase I. Nature 502, 644–649 (2013).

7. Engel, C., Sainsbury, S., Cheung, A. C., Kostrewa, D. & Cramer, P. RNA polymerase I structure and transcription regulation. Nature 502, 650–655 (2013).

8. Hoffmann, N. A. et al. Molecular structures of unbound and transcribing RNA polymerase III. Nature 528, 231–236 (2015).

9. Gietl, A. et al. Eukaryotic and archaeal TBP and TFB/TF(II)B follow different promoter DNA bending pathways. Nucleic Acids Research 42, 6219–6231 (2014).

10. Werner, F. & Weinzierl, R. O. J. Direct Modulation of RNA Polymerase Core Functions by Basal Transcription Factors. Molecular and Cellular Biology 25, 8344–8355 (2005).

11. Smollett, K., Blombach, F., Reichelt, R., Thomm, M. & Werner, F. A global analysis of transcription reveals two modes of Spt4/5 recruitment to archaeal RNA polymerase. Nat Microbiol 2, 17021 (2017).

12. Blombach, F., Fouqueau, T., Matelska, D., Smollett, K. & Werner, F. Promoter-proximal elongation regulates transcription in archaea. Nat Commun 12, 5524 (2021).

13. Soppa, J. Transcription initiation in Archaea: facts, factors and future aspects. Mol Microbiol 31, 1295–1305 (1999).

14. Ouhammouch, M., Dewhurst, R. E., Hausner, W., Thomm, M. & Geiduschek, E. P. Activation of archaeal transcription by recruitment of the TATA-binding protein. Proc. Natl. Acad. Sci. U.S.A. 100, 5097–5102 (2003).

15. Bell, S. D., Kosa, P. L., Sigler, P. B. & Jackson, S. P. Orientation of the transcription preinitiation complex in Archaea. Proc. Natl. Acad. Sci. U.S.A. 96, 13662–13667 (1999).

16. Nagy, J. et al. Complete architecture of the archaeal RNA polymerase open complex from single-molecule FRET and NPS. Nat Commun 6, 6161 (2015).

17. Grohmann, D. et al. The Initiation Factor TFE and the Elongation Factor Spt4/5 Compete for the RNAP Clamp during Transcription Initiation and Elongation. Molecular Cell 43, 263–274 (2011).

18. Blombach, F. et al. Archaeal TFEα/β is a hybrid of TFIIE and the RNA polymerase III subcomplex hRPC62/39. eLife 4, e08378 (2015).

19. Schulz, S. et al. TFE and Spt4/5 open and close the RNA polymerase clamp during the transcription cycle. Proc. Natl. Acad. Sci. U.S.A. 113, (2016).

20. Hirtreiter, A. et al. Spt4/5 stimulates transcription elongation through the RNA polymerase clamp coiled-coil motif. Nucleic Acids Research 38, 4040–4051 (2010).

21. Martinez-Rucobo, F. W., Sainsbury, S., Cheung, A. C. & Cramer, P. Architecture of the RNA polymerase-Spt4/5 complex and basis of universal transcription processivity: RNA polymerase-Spt4/5 complex architecture. The EMBO Journal 30, 1302–1310 (2011).

22. Klein, B. J. et al. RNA polymerase and transcription elongation factor Spt4/5 complex structure. Proc. Natl. Acad. Sci. U.S.A. 108, 546–550 (2011).

23. Werner, F. A Nexus for Gene Expression—Molecular Mechanisms of Spt5 and NusG in the Three Domains of Life. Journal of Molecular Biology 417, 13–27 (2012).

24. Decker, T.-M. Mechanisms of Transcription Elongation Factor DSIF (Spt4–Spt5). Journal of Molecular Biology 433, 166657 (2021).

25. Sanders, T. J. et al. TFS and Spt4/5 accelerate transcription through archaeal histone-based chromatin: Transcription through archaeal chromatin. Mol Microbiol 111, 784–797 (2019).

26. Landick, R. Transcriptional Pausing as a Mediator of Bacterial Gene Regulation. Annu. Rev. Microbiol. 75, 291–314 (2021).

27. Lawson, M. R. et al. Mechanism for the Regulated Control of Bacterial Transcription Termination by a Universal Adaptor Protein. Molecular Cell 71, 911–922.e4 (2018).

28. Wang, C. et al. Structural basis of transcription-translation coupling. Science 369, 1359–1365 (2020).

29. Weixlbaumer, A., Grünberger, F., Werner, F. & Grohmann, D. Coupling of Transcription and Translation in Archaea: Cues From the Bacterial World. Front. Microbiol. 12, 661827 (2021).

30. Webster, M. W. et al. Structural basis of transcription-translation coupling and collision in bacteria. Science 369, 1355–1359 (2020).

31. Blaha, G. M. & Wade, J. T. Transcription-Translation Coupling in Bacteria. Annu. Rev. Genet. 56, 187–205 (2022).

32. Yakhnin, A. V. et al. NusG controls transcription pausing and RNA polymerase translocation throughout the *Bacillus subtilis* genome. Proc. Natl. Acad. Sci. U.S.A. 117, 21628–21636 (2020).

33. Nedialkov, Y., Svetlov, D., Belogurov, G. A. & Artsimovitch, I. Locking the nontemplate DNA to control transcription: RfaH remodels DNA in the transcription elongation complex. Molecular Microbiology 109, 445–457 (2018).

34. Kang, J. Y. et al. Structural Basis for Transcript Elongation Control by NusG Family Universal Regulators. Cell 173, 1650–1662.e14 (2018).

35. Vishwakarma, R. K., Qayyum, M. Z., Babitzke, P. & Murakami, K. S. Allosteric mechanism of transcription inhibition by NusG-dependent pausing of RNA polymerase. Proc. Natl. Acad. Sci. U.S.A. 120, e2218516120 (2023).

36. Engel, C., Neyer, S. & Cramer, P. Distinct Mechanisms of Transcription Initiation by RNA Polymerases I and II. Annu. Rev. Biophys. 47, 425–446 (2018).

37. Sanders, T. J. et al. FttA is a CPSF73 homologue that terminates transcription in Archaea. Nat Microbiol 5, 545–553 (2020).

38. Marshall, C. J., Qayyum, M. Z., Walker, J. E., Murakami, K. S. & Santangelo, T. J. The structure and activities of the archaeal transcription termination factor Eta detail vulnerabilities of the transcription elongation complex. Proc. Natl. Acad. Sci. U.S.A. 119, e2207581119 (2022).

39. Girbig, M., Misiaszek, A. D. & Müller, C. W. Structural insights into nuclear transcription by eukaryotic DNA-dependent RNA polymerases. Nat Rev Mol Cell Biol 23, 603–622 (2022).

40. Wolberger, C. How structural biology transformed studies of transcription regulation. Journal of Biological Chemistry 296, 100741 (2021).

41. Mohamed, A. A., Vazquez Nunez, R. & Vos, S. M. Structural advances in transcription elongation. Current Opinion in Structural Biology 75, 102422 (2022).

42. Nogales, E., Louder, R. K. & He, Y. Structural Insights into the Eukaryotic Transcription Initiation Machinery. Annu. Rev. Biophys. 46, 59–83 (2017).

43. Pilsl, M. & Engel, C. Structural Studies of Eukaryotic RNA Polymerase I Using Cryo-Electron Microscopy. in Ribosome Biogenesis (ed. Entian, K.-D.) vol. 2533 71–80 (Springer US, 2022).

44. Kusser, A. G. et al. Structure of an Archaeal RNA Polymerase. Journal of Molecular Biology 376, 303–307 (2008).

45. Jun, S.-H. et al. The X-ray crystal structure of the euryarchaeal RNA polymerase in an open-clamp configuration. Nat Commun 5, 5132 (2014).

46. Korkhin, Y. et al. Evolution of Complex RNA Polymerases: The Complete Archaeal RNA Polymerase Structure. PLoS Biol 7, e1000102 (2009).

47. Hirata, A., Klein, B. J. & Murakami, K. S. The X-ray crystal structure of RNA polymerase from Archaea. Nature 451, 851–854 (2008).

48. Jun, S.-H. et al. Direct binding of TFEα opens DNA binding cleft of RNA polymerase. Nat Commun 11, 6123 (2020).

49. Sheppard, C. et al. Repression of RNA polymerase by the archaeo-viral regulator ORF145/RIP. Nat Commun 7, 13595 (2016).

50. Wojtas, M. N., Mogni, M., Millet, O., Bell, S. D. & Abrescia, N. G. A. Structural and functional analyses of the interaction of archaeal RNA polymerase with DNA. Nucleic Acids Research 40, 9941–9952 (2012).

51. Fouqueau, T. et al. The transcript cleavage factor paralogue TFS4 is a potent RNA polymerase inhibitor. Nat Commun 8, 1914 (2017).

52. Punjani, A., Rubinstein, J. L., Fleet, D. J. & Brubaker, M. A. cryoSPARC: algorithms for rapid unsupervised cryo-EM structure determination. Nat Methods 14, 290–296 (2017).

53. Neyer, S. et al. Structure of RNA polymerase I transcribing ribosomal DNA genes. Nature 540, 607–610 (2016).

54. Bernecky, C., Herzog, F., Baumeister, W., Plitzko, J. M. & Cramer, P. Structure of transcribing mammalian RNA polymerase II. Nature 529, 551–554 (2016).

55. Ehara, H. et al. Structure of the complete elongation complex of RNA polymerase II with basal factors. Science 357, 921–924 (2017).

56. Bernecky, C., Plitzko, J. M. & Cramer, P. Structure of a transcribing RNA polymerase II–DSIF complex reveals a multidentate DNA–RNA clamp. Nat Struct Mol Biol 24, 809–815 (2017).

57. Wang, C., et al. Structural basis of archaeal FttA-dependent transcription termination. http://biorxiv.org/lookup/doi/10.1101/2023.08.09.552649 (2023) doi:10.1101/2023.08.09.552649.

58. Weixlbaumer, A., Leon, K., Landick, R. & Darst, S. A. Structural Basis of Transcriptional Pausing in Bacteria. Cell 152, 431–441 (2013).

59. Landick, R. RNA Polymerase Clamps Down. Cell 105, 567–570 (2001).

60. Lin, W. et al. Structural Basis of Transcription Inhibition by Fidaxomicin (Lipiarmycin A3). Molecular Cell 70, 60–71.e15 (2018).

61. Finn, R. D. Escherichia coli RNA polymerase core and holoenzyme structures. The EMBO Journal 19, 6833–6844 (2000).

62. Duchi, D., Mazumder, A., Malinen, A. M., Ebright, R. H. & Kapanidis, A. N. The RNA polymerase clamp interconverts dynamically among three states and is stabilized in a partly closed state by ppGpp. Nucleic Acids Research 46, 7284–7295 (2018).

63. Qayyum, M. Z., Molodtsov, V., Renda, A. & Murakami, K. S. Structural basis of RNA polymerase recycling by the Swi2/Snf2 family of ATPase RapA in Escherichia coli. Journal of Biological Chemistry 297, 101404 (2021).

64. Bellecourt, M. J., Ray-Soni, A., Harwig, A., Mooney, R. A. & Landick, R. RNA Polymerase Clamp Movement Aids Dissociation from DNA but Is Not Required for RNA Release at Intrinsic Terminators. Journal of Molecular Biology 431, 696–713 (2019).

65. Chakraborty, A. et al. Opening and Closing of the Bacterial RNA Polymerase Clamp. Science 337, 591–595 (2012).

66. Pilotto, S. et al. Structural basis of RNA polymerase inhibition by viral and host factors. Nat Commun 12, 5523 (2021).

67. Fianu, I., Dienemann, C., Aibara, S., Schilbach, S. & Cramer, P. Cryo-EM structure of mammalian RNA polymerase II in complex with human RPAP2. Commun Biol 4, 606 (2021).

68. Aibara, S., Dienemann, C. & Cramer, P. Structure of an inactive RNA polymerase II dimer. Nucleic Acids Research 49, 10747–10755 (2021).

69. Engel, C. et al. Structural Basis of RNA Polymerase I Transcription Initiation. Cell 169, 120–131.e22 (2017).

70. Tafur, L. et al. Molecular Structures of Transcribing RNA Polymerase I. Molecular Cell 64, 1135– 1143 (2016).

71. Neyer, S. et al. Structure of RNA polymerase I transcribing ribosomal DNA genes. Nature 540, 607–610 (2016).

72. Heiss, F. B., Daiß, J. L., Becker, P. & Engel, C. Conserved strategies of RNA polymerase I hibernation and activation. Nat Commun 12, 758 (2021).

73. Daiß, J. L. et al. The human RNA polymerase I structure reveals an HMG-like docking domain specific to metazoans. Life Sci. Alliance 5, e202201568 (2022).

74. Liu, B. & Steitz, T. A. Structural insights into NusG regulating transcription elongation. Nucleic Acids Res 45, 968–974 (2017).

75. Wang, Q., Daiß, J. L., Xu, Y. & Engel, C. Snapshots of RNA polymerase III in action – A mini review. Gene 821, 146282 (2022).

76. Blombach, F., Smollett, K. L., Grohmann, D. & Werner, F. Molecular Mechanisms of Transcription Initiation—Structure, Function, and Evolution of TFE/TFIIE-Like Factors and Open Complex Formation. Journal of Molecular Biology 428, 2592–2606 (2016).

77. Blombach, F. et al. Archaeology of RNA polymerase: factor swapping during the transcription cycle. Biochemical Society Transactions 41, 362–367 (2013).

78. Waege, I., Schmid, G., Thumann, S., Thomm, M. & Hausner, W. Shuttle Vector-Based Transformation System for *Pyrococcus furiosus*. Appl Environ Microbiol 76, 3308–3313 (2010).

79. Sommer, B. et al. Activation of a Chimeric Rpb5/RpoH Subunit Using Library Selection. PLoS ONE 9, e87485 (2014).

80. Fiala, G. & Stetter, K. O. Pyrococcus furiosus sp. nov. represents a novel genus of marine heterotrophic archaebacteria growing optimally at 100°C. Arch. Microbiol. 145, 56–61 (1986).

81. Gibson, D. G. Enzymatic Assembly of Overlapping DNA Fragments. in Methods in Enzymology vol. 498 349–361 (Elsevier, 2011).

82. Department of Computational Biology & Bioinformatics, Jacob School of Biotechnology & Bioengineering, Sam Higginbottom Institute of Agriculture Technology & Sciences, Allahabad-211007, Uttar Pradesh, Bharat (India) et al. MFPPI – Multi FASTA ProtParam Interface. Bioinformation 12, 74–77 (2016).

83. Pilsl, M. et al. Preparation of RNA Polymerase Complexes for Their Analysis by Single-Particle Cryo-Electron Microscopy. in Ribosome Biogenesis (ed. Entian, K.-D.) vol. 2533 81–96 (Springer US, 2022).

84. Schorb, M., Haberbosch, I., Hagen, W. J. H., Schwab, Y. & Mastronarde, D. N. Software tools for automated transmission electron microscopy. Nat Methods 16, 471–477 (2019).

85. Alegria-Schaffer, A. General Protein–Protein Cross-Linking. in Methods in Enzymology vol. 539 81–87 (Elsevier, 2014).

86. Scheres, S. H. W. Amyloid structure determination in *RELION* -3.1. Acta Crystallogr D Struct Biol 76, 94–101 (2020).

87. Zheng, S. Q. et al. MotionCor2: anisotropic correction of beam-induced motion for improved cryo-electron microscopy. Nat Methods 14, 331–332 (2017).

88. Bepler, T. et al. Positive-unlabeled convolutional neural networks for particle picking in cryo-electron micrographs. Nat Methods 16, 1153–1160 (2019).

89. Tan, Y. Z. et al. Addressing preferred specimen orientation in single-particle cryo-EM through tilting. Nat Methods 14, 793–796 (2017).

90. Wagner, T. et al. SPHIRE-crYOLO is a fast and accurate fully automated particle picker for cryo-EM. Commun Biol 2, 218 (2019).

91. Jakobi, A. J., Wilmanns, M. & Sachse, C. Model-based local density sharpening of cryo-EM maps. eLife 6, e27131 (2017).

92. (IUCr) Recent developments in the CCP-EMsoftware suite. http://scripts.iucr.org/cgi-bin/paper?S2059798317007859.

93. Waterhouse, A. et al. SWISS-MODEL: homology modelling of protein structures and complexes. Nucleic Acids Research 46, W296–W303 (2018).

94. Emsley, P. & Cowtan, K. *Coot* : model-building tools for molecular graphics. Acta Crystallogr D Biol Crystallogr 60, 2126–2132 (2004).

95. Croll, T. I. *ISOLDE* : a physically realistic environment for model building into low-resolution electron-density maps. Acta Crystallogr D Struct Biol 74, 519–530 (2018).

96. Pettersen, E. F. et al. UCSF CHIMERAX : Structure visualization for researchers, educators, and developers. Protein Science 30, 70–82 (2021).

97. Adams, P. D., et al. *PHENIX* : a comprehensive Python-based system for macromolecular structure solution. Acta Crystallogr D Biol Crystallogr 66, 213–221 (2010).

98. Atanassov, I. & Urlaub, H. Increased proteome coverage by combining PAGE and peptide isoelectric focusing: Comparative study of gel-based separation approaches. Proteomics 13, 2947–2955 (2013).

