## Supplemental Tables and Figures for "Structural basis of archaeal RNA polymerase transcription elongation and Spt4/5 recruitment"

**Figure S1**

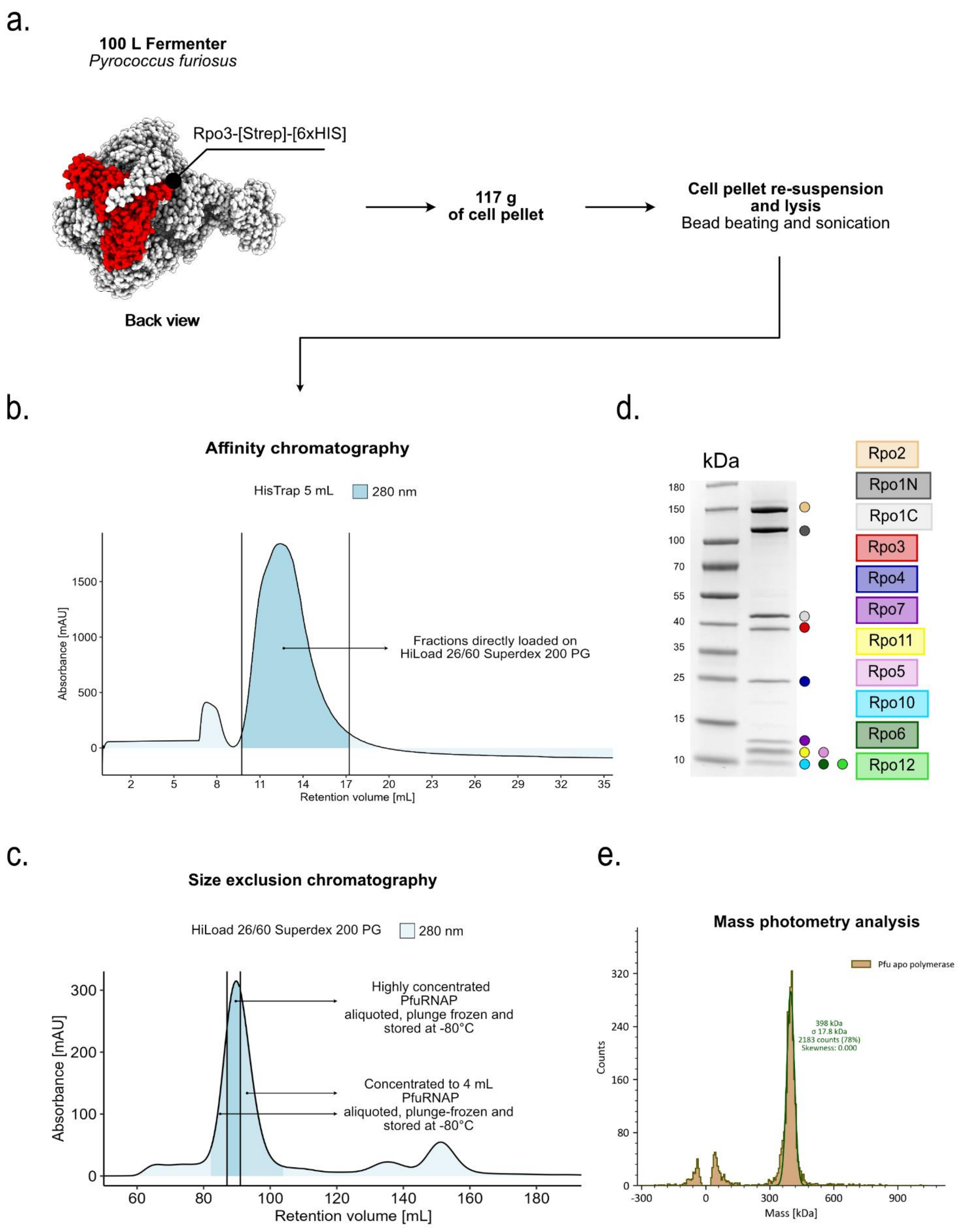
a.
100 L Fermenter
Pyrococcus furiosus
Rpo3-[Strep]-[6xHIS]
Back view
117 g
of cell pellet
Cell pellet re-suspension
and lysis
Bead beating and sonication
b.
Affinity chromatography
HisTrap 5 mL
280 nm
Absorbance [mAU]
1500
1000
500
0
Fractions directly loaded on
HiLoad 26/60 Superdex 200 PG
2 5 8 11 14 17 20 23 26 29 32 35
Retention volume [mL]
c.
Size exclusion chromatography
HiLoad 26/60 Superdex 200 PG
280 nm
Absorbance [mAU]
300
200
100
0
Highly concentrated
PfuRNAP
aliquoted, plunge frozen and
stored at -80°C
Concentrated to 4 mL
PfuRNAP
aliquoted, plunge-frozen and
stored at -80°C
60 80 100 120 140 160 180
Retention volume [mL]
d.
kDa
180
150
100
70
55
40
35
25
15
10
Rpo2
Rpo1N
Rpo1C
Rpo3
Rpo4
Rpo7
Rpo11
Rpo5
Rpo10
Rpo6
Rpo12
e.
Mass photometry analysis
Pfu apo polymerase
398 kDa
σ 17.8 kDa
2183 counts (78%)
Skewness: 0.000
Counts
320
240
160
80
0
-300 0 300 600 900
Mass [kDa]

**Figure S1. Schematic representation of *Pyrococcus furiosus* endogenous RNA polymerase purification. a.** *Pfu* RNAP was purified from a genetically engineered strain that expresses a genomic copy of Rpo3 (shown in red) genetically fused to a Strep- and 6xHIS-tag. Large scale cultivation of this strain in a 100 L fermenter resulted in 117 g of cell pellet. **b.** RNAP was purified from cell lysate via Ni-NTA affinity chromatography exploiting the 6xHIS-tag. **c.** Finally, size exclusion chromatography was performed with the most concentrated fractions of the affinity chromatography. **d.** SDS-PAGE analysis of the peak fraction from the size exclusion chromatography using a polyacrylamide gradient (gradient concentration 4%-20%) shows bands for all RNAP subunits indicating successful purification. **e.** Mass photometry analysis of the purified *pfu* RNAP displaying a peak at the expected mass.

**Figure S2**

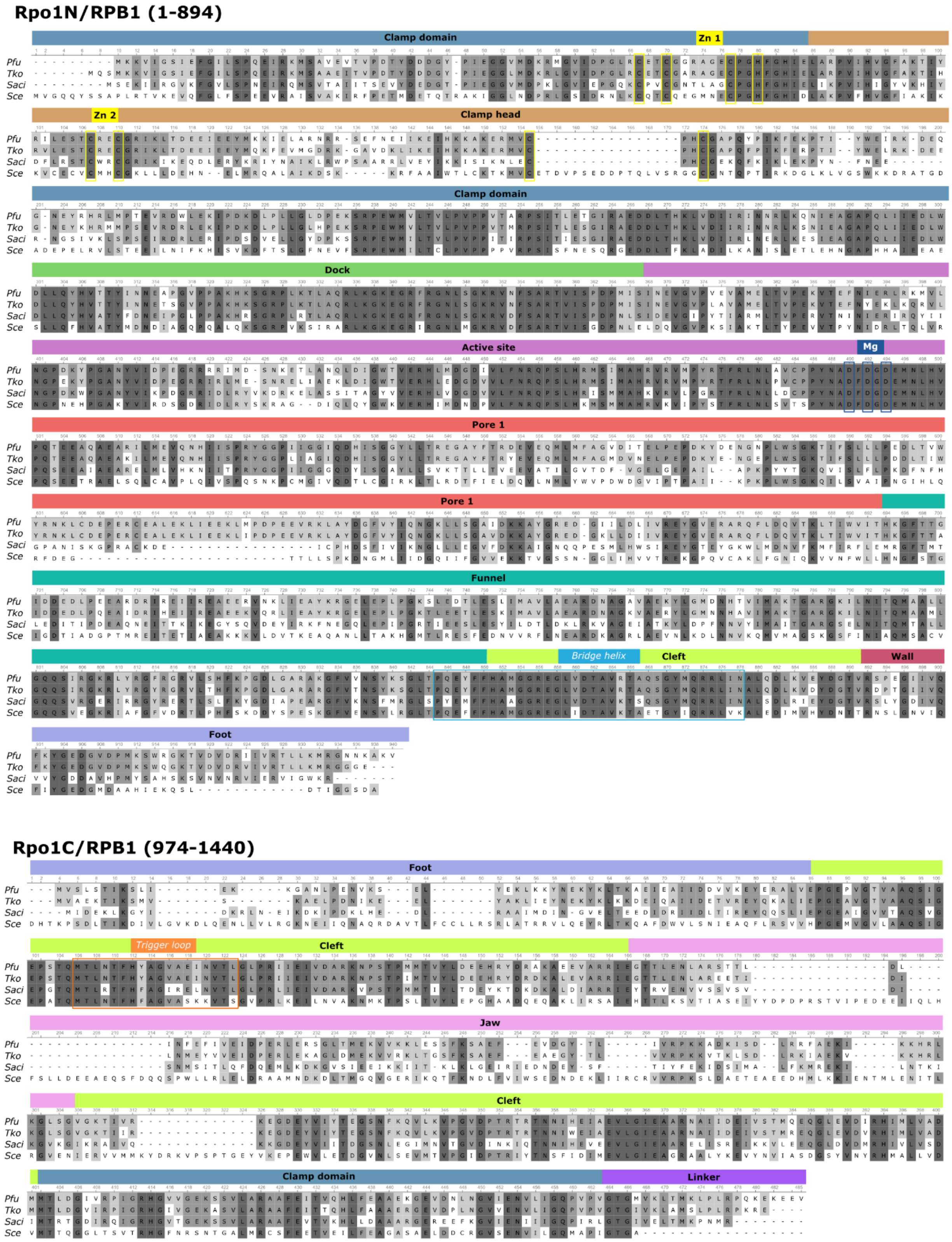
Rpo1N/RPB1 (1-894)
Clamp domain
Zn 1
Zn 2
Clamp head
Clamp domain
Dock
Active site
Mg
Pore 1
Pore 1
Funnel
Bridge helix
Cleft
Wall
Foot
Pfu
Tko
Saci
Sce
Rpo1C/RPB1 (974-1440)
Foot
Trigger loop
Cleft
Jaw
Cleft
Clamp domain
Linker

### Rpo2/RPB2

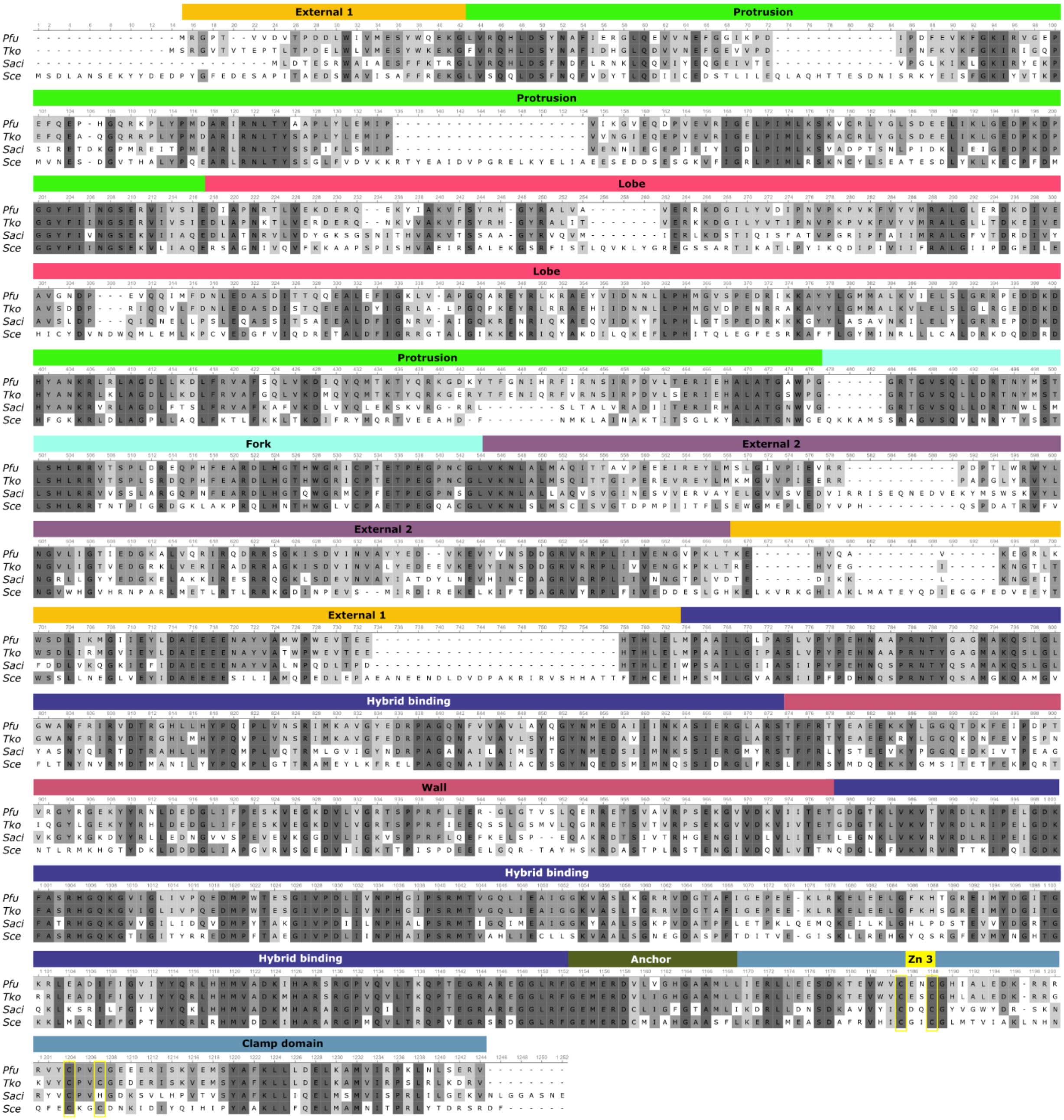

External 1
Protrusion
Lobe
Fork
External 2
Hybrid binding
Wall
Anchor
Zn 3
Clamp domain
Pfu
Tko
Saci
Sce

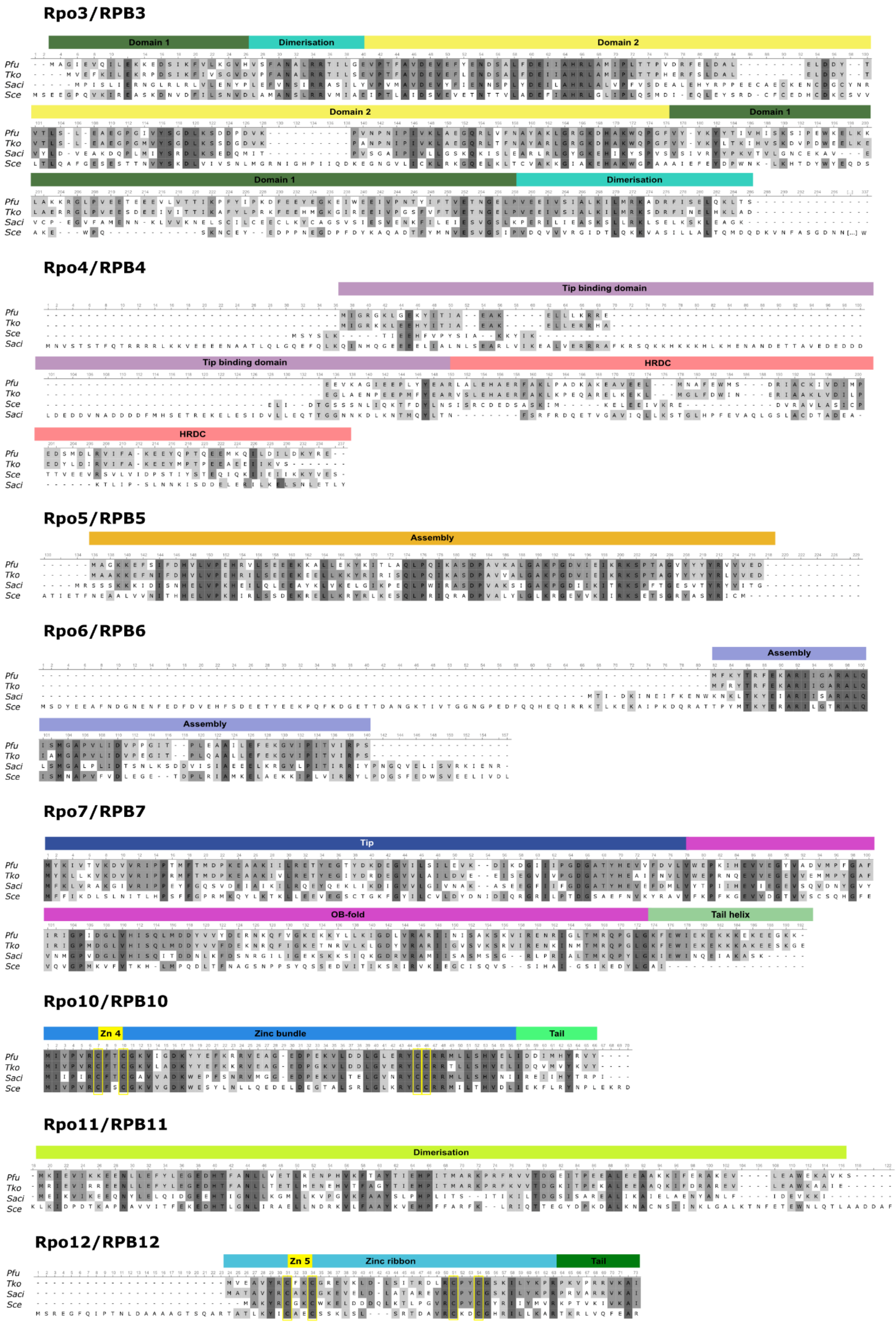
Rpo3/RPB3
Domain 1
Dimerisation
Domain 2
Domain 2
Domain 1
Domain 1
Dimerisation
Pfu
Tko
Saci
Sce
Rpo4/RPB4
Tip binding domain
Tip binding domain
HRDC
HRDC
Pfu
Tko
Sce
Saci
Rpo5/RPB5
Assembly
Pfu
Tko
Saci
Sce
Rpo6/RPB6
Assembly
Assembly
Rpo7/RPB7
Tip
OB-fold
Tail helix
Rpo10/RPB10
Zn 4
Zinc bundle
Tail
Rpo11/RPB11
Dimerisation
Rpo12/RPB12
Zn 5
Zinc ribbon
Tail

**Figure S2. Structure-based sequence alignment of archaeal and eukaryotic RNAP subunits.** Structure-based sequence alignment of *Pfu* RNAP subunits with the respective counterparts from *Thermococcus kodakarensis* (pdb 6KF3)[1], *Sulfolobus acidocaldarius* (pdb 7OK0)[2], and *Saccharomyces cerevisieae* (pdb 6GYK)[3]. The sequence alignment was performed with ClustalO and the figures were produced and modified with Unipro UGENE and Affinity Designer 2.1.1. Domain assignment was performed according to the 3D structure of *S. acidocaldarius*. Zinc and Magnesium coordinating amino acids are marked in yellow and dark blue, respectively.

### Figure S3

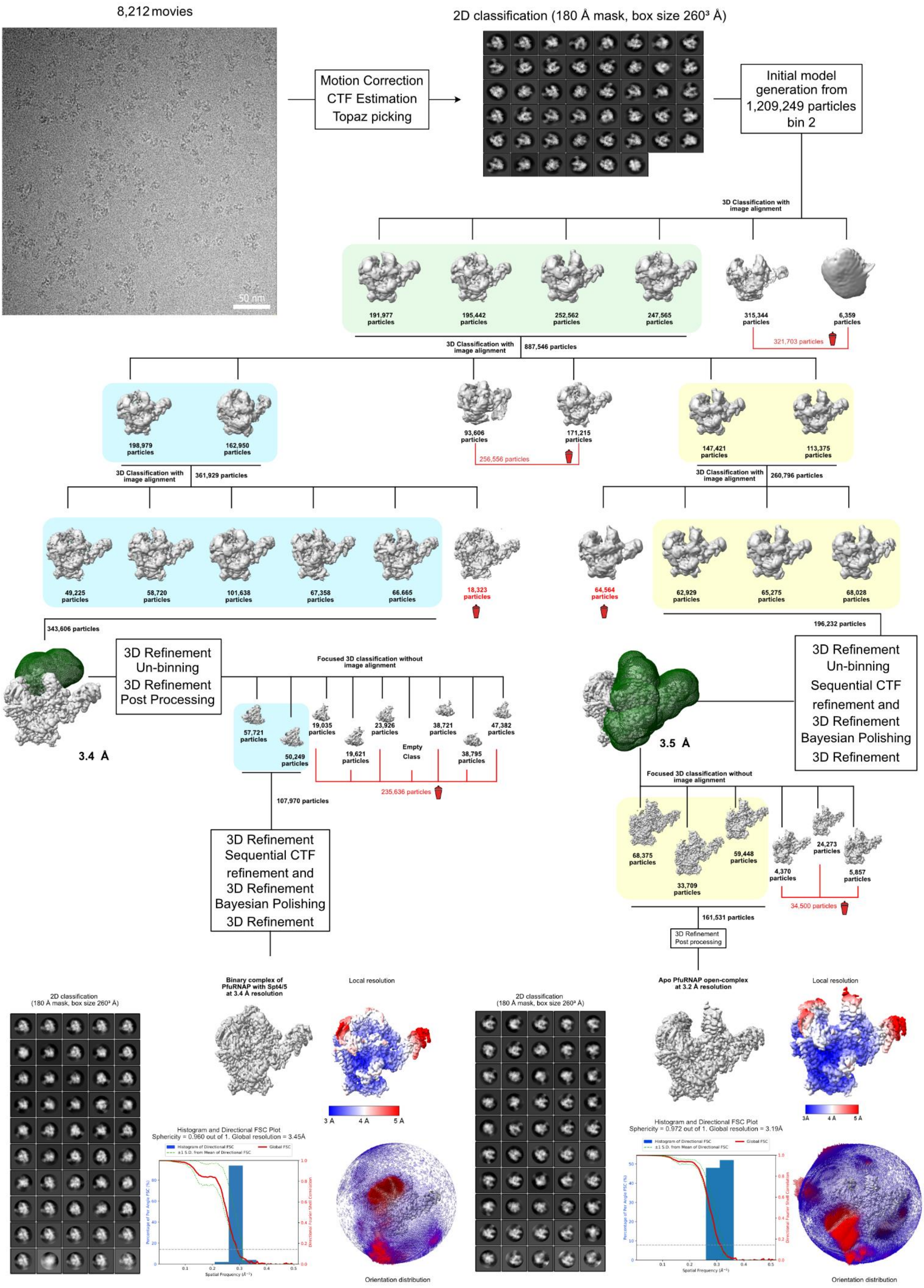
8,212 movies
50 nm
Motion Correction
CTF Estimation
Topaz picking
2D classification (180 Å mask, box size 260³ Å)
Initial model generation from 1,209,249 particles bin 2
3D Classification with image alignment
191,977 particles
195,442 particles
252,562 particles
247,565 particles
315,344 particles
6,359 particles
321,703 particles
3D Classification with image alignment
887,546 particles
198,979 particles
162,950 particles
93,606 particles
171,215 particles
256,556 particles
147,421 particles
113,375 particles
3D Classification with image alignment
361,929 particles
3D Classification with image alignment
260,796 particles
49,225 particles
58,720 particles
101,638 particles
67,358 particles
66,665 particles
18,323 particles
64,564 particles
62,929 particles
65,275 particles
68,028 particles
343,606 particles
196,232 particles
3D Refinement
Un-binning
3D Refinement
Post Processing
3.4 Å
Focused 3D classification without image alignment
57,721 particles
50,249 particles
19,035 particles
19,621 particles
23,926 particles
Empty Class
38,721 particles
38,795 particles
47,382 particles
235,636 particles
107,970 particles
3D Refinement
Sequential CTF refinement and
3D Refinement
Bayesian Polishing
3D Refinement
3D Refinement
Un-binning
Sequential CTF refinement and
3D Refinement
Bayesian Polishing
3D Refinement
3.5 Å
Focused 3D classification without image alignment
68,375 particles
33,709 particles
59,448 particles
4,370 particles
24,273 particles
5,857 particles
34,500 particles
161,531 particles
3D Refinement
Post processing
2D classification
(180 Å mask, box size 260³ Å)
Binary complex of PfuRNAP with Spt4/5 at 3.4 Å resolution
Local resolution
3 Å
4 Å
5 Å
Histogram and Directional FSC Plot
Sphericity = 0.960 out of 1. Global resolution = 3.45Å
Orientation distribution
2D classification
(180 Å mask, box size 260³ Å)
Apo PfuRNAP open-complex at 3.2 Å resolution
Local resolution
3Å
4Å
5Å
Histogram and Directional FSC Plot
Sphericity = 0.972 out of 1. Global resolution = 3.19Å
Orientation distribution

2

**Figure S3. Cryo-EM and single particle analysis processing workflow for the apo *Pfu* RNAP sample, including representative micrograph, 2D averages, 3D classifications, final refined map, local resolution heatmap, Fourier completeness plot, histogram and directional FSC in Relion 4.0.**

**Figure S4**

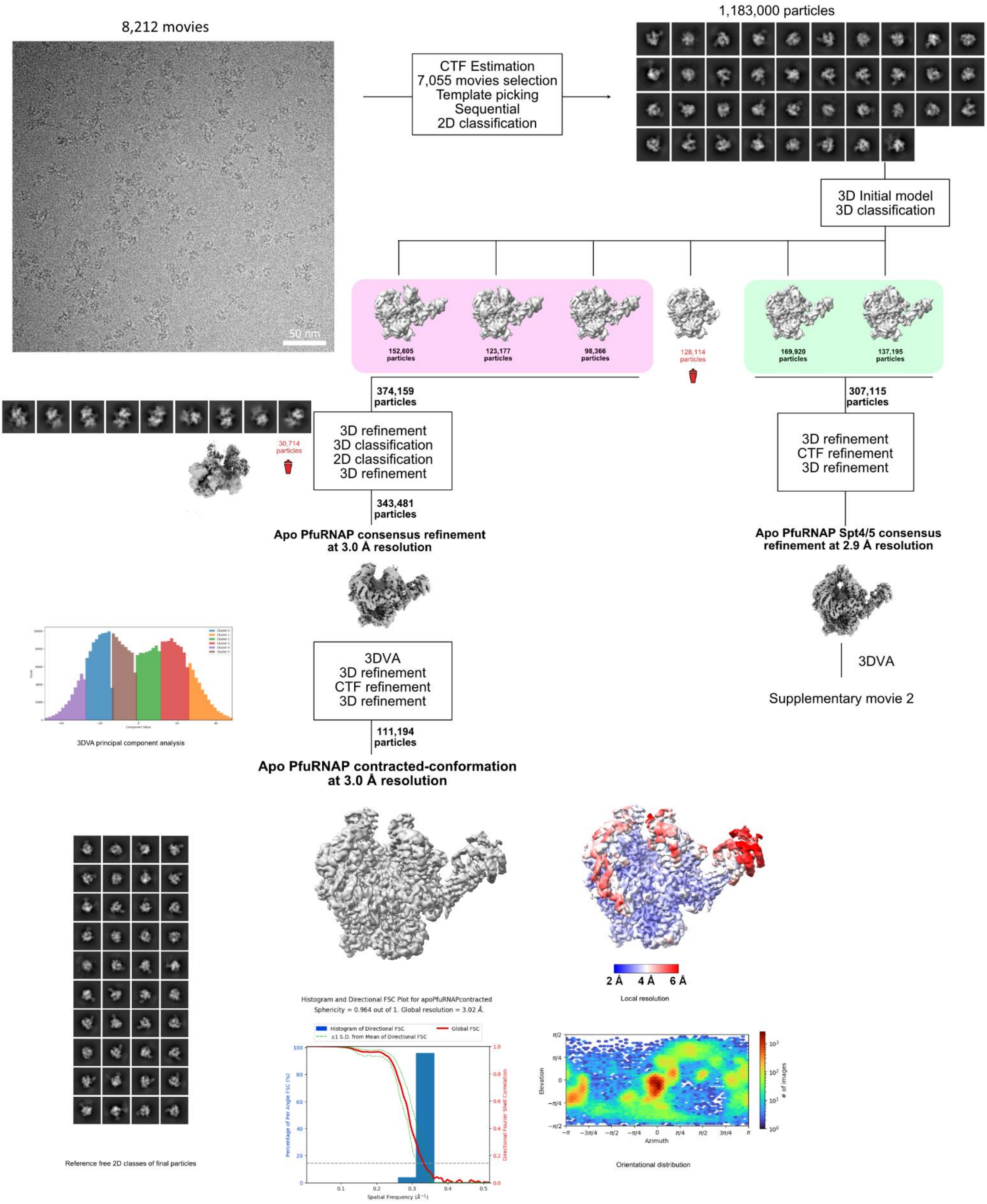


**Figure S4. Cryo-EM and single particle analysis processing workflow for the apo PfuRNAP sample in cryosparc, including representative micrograph; 2D averages; 3D classifications, intermediate and final refined map; 3DVA cluster and analysis, local resolution heatmap; Fourier completeness plot; histogram and directional FSC;**

**Figure S5**

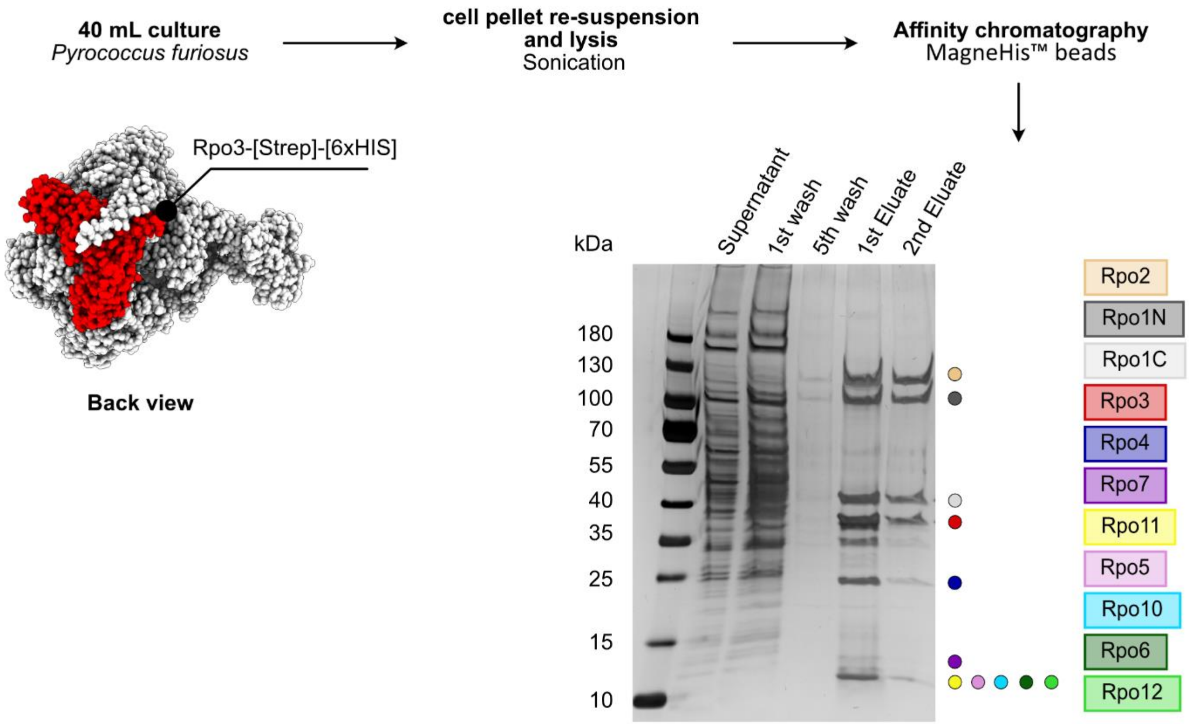


**Figure S5. Quality control sample preparation for mass spectrometry.** SDS-PAGE analysis of *Pfu* RNAP purification in small scale using MagneHIS$^{TM}$ protein purification system for mass spectrometry analysis. Shown are the indicated purifications steps which were analyzed on a 4%-20% gradient gel.

### Figure S6

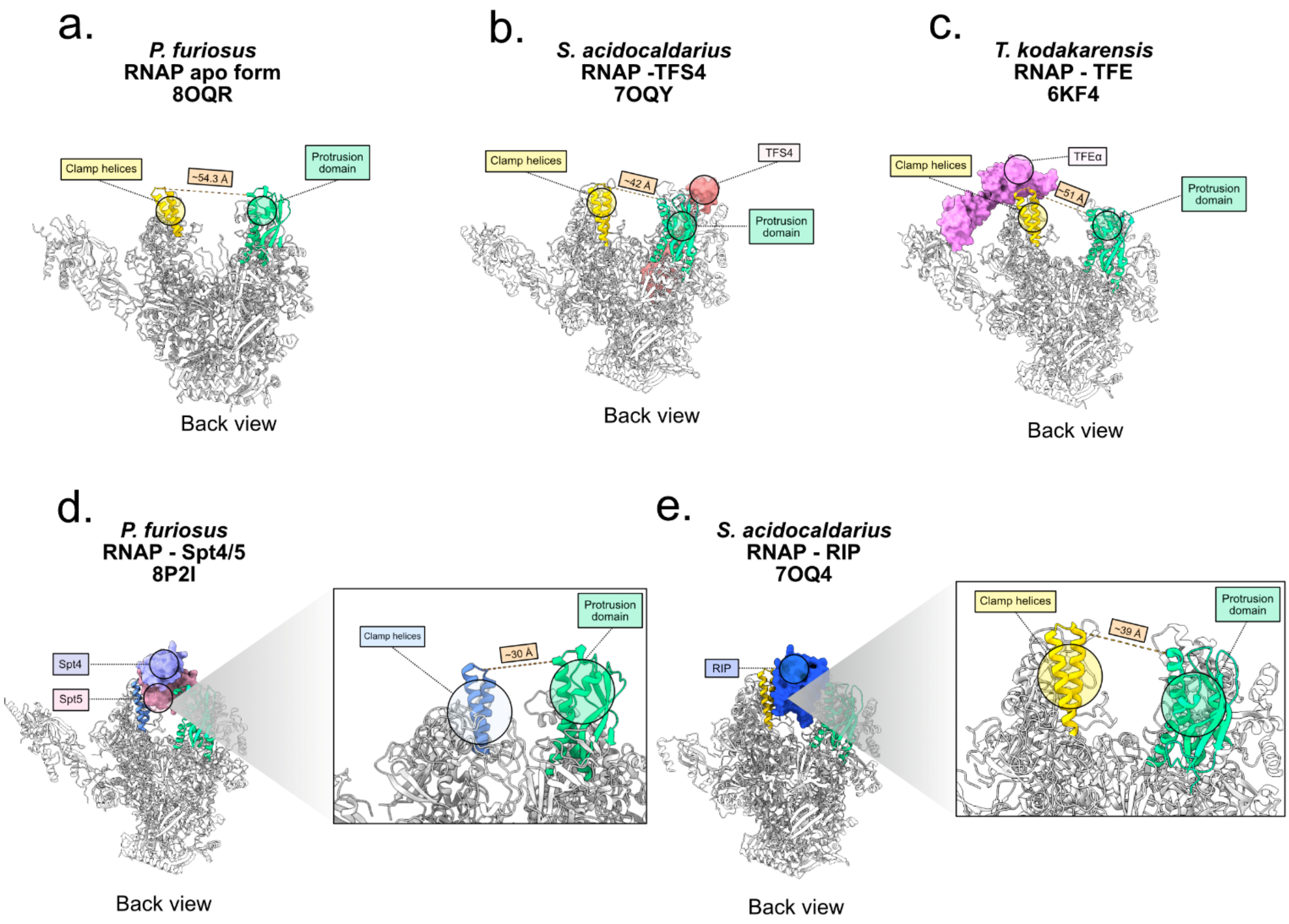


**Figure S6. Conformation of the clamp domain across archaeal RNAPs and transcription complexes**. **a.** In the apo state of *Pfu* RNAP, the clamp adopts an open clamp state with ~54.3 Å between Rpo1N-G$^{259}$ in the coiled-coil motif on the clamp domain (yellow) and Rpo2-R$^{377}$ in the protrusion domain (green). **b.** *Sulfolobus acidocaldarius* RNAP in complex with TFS4[2] is characterized by a closed clamp domain with a distance of ~42 Å between residues Rpo1N-G$^{257}$ and Rpo2-R$^{373}$. **c.** *Thermococcus kodakarensis* TFEα[1] shifts the clamp domain to an open state with a width between the coiled-coil motif (Rpo1N-A$^{257}$) and protrusion (Rpo2-R$^{380}$) of ~51Å. **d.** Spt4/5 binding to the *Pfu* RNAP in absence of nucleic acids leads to a closed clamp conformation (blue). The distance between clamp (Rpo1N-G$^{259}$) and protrusion domain (Rpo2-R$^{377}$) is ~30 Å. **e.** The protein RIP[2] is of viral origin and binds the RNAP from *Sulfolobus acidocaldarius.* Similar to Spt4/5, RIP binds inside the DNA binding channel. However, the distance between clamp (Rpo1N-G$^{257}$) and protrusion (Rpo2-R$^{373}$) differs by about ~9 Å with a more open clamp in case of the RNAP-RIP complex.

**Figure S7**

a. **PfuRNAP cryo-EM sample preparation**

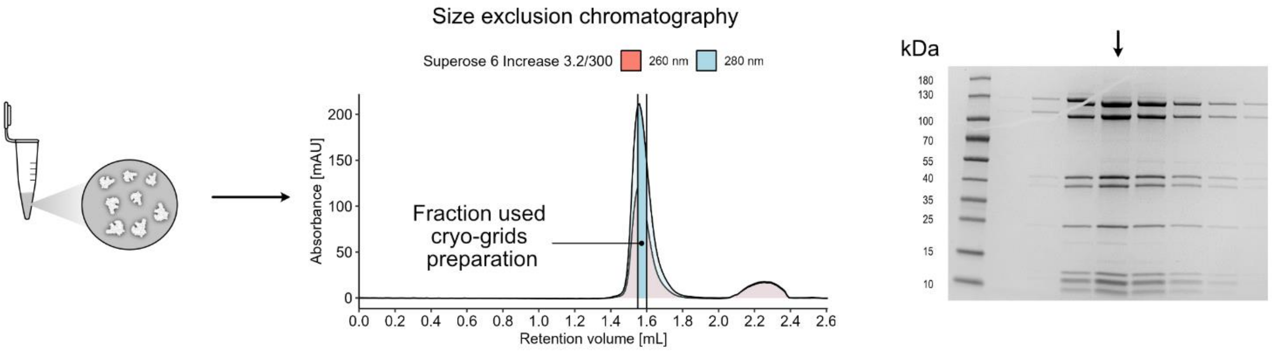


b. **Negative staining analysis**

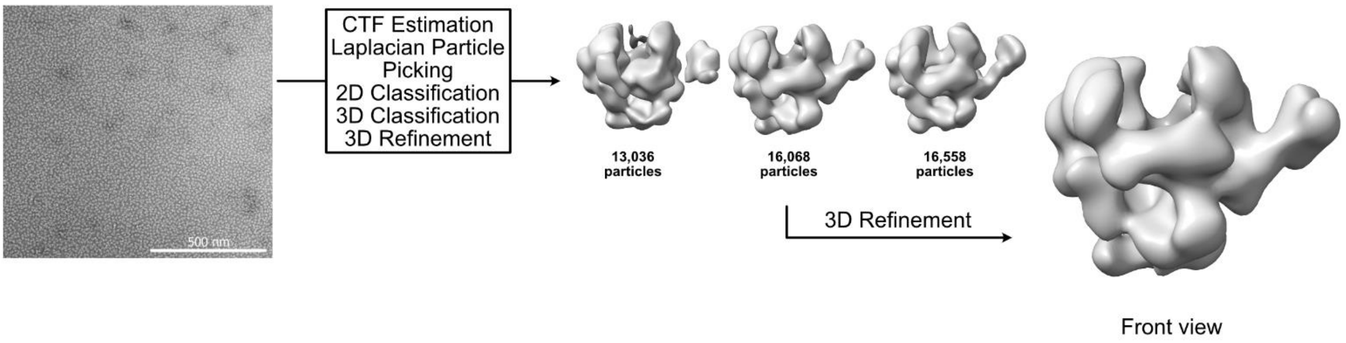


c. **TEC assembly for cryo-EM**

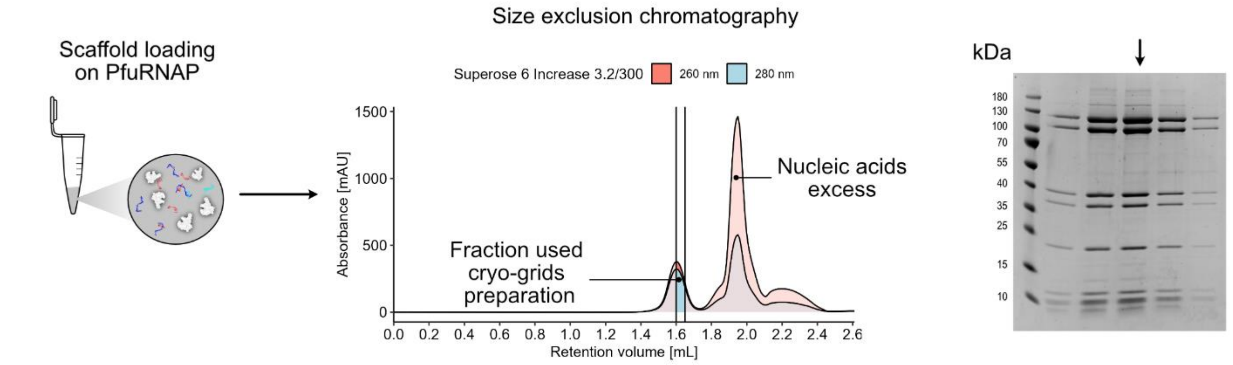


d. **Negative staining analysis**

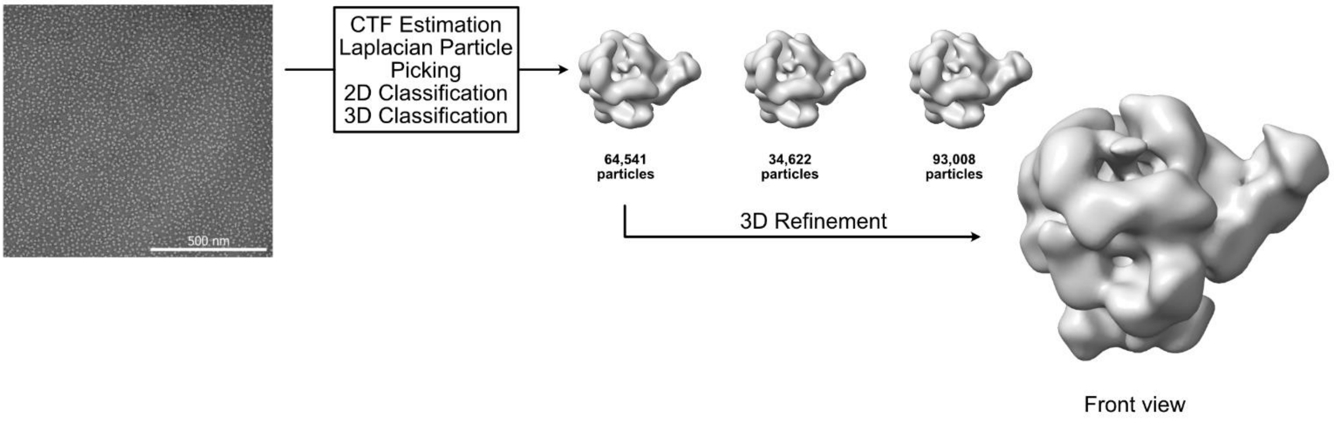

**Figure S7. Sample preparation and negative staining for the apo enzyme of *Pfu* RNAP and transcription elongation complex samples. a.** Chromatogram of the size exclusion chromatography run of apo *Pfu* RNAP shows a homogeneous elution with dominant absorbance at 280 nm (light blue). Successful purification of *Pfu* RNAP was confirmed by SDA-PAGE using a gradient gel (4%-20%). **b.** Negative staining of apo *Pfu* RNAP. A small dataset was acquired (23 micrographs) yielding a low-resolution density map of *Pfu* RNAP with an open-clamp conformation. **c.** Sample preparation of the transcription elongation complex (TEC) containing *Pfu* RNAP and a synthetic elongation scaffold. After incubation, size exclusion chromatography was performed to separate the TEC from free nucleic acids. The chromatogram indicates successful formation of the TEC as absorbance at 260 nm (salmon) is higher than at 280 nm in the polymerase peak. The success of the purification was confirmed by 4%-20% gradient SDS-PAGE. **d.** A dataset of negatively stained particles was collected (161 micrographs) leading to a reconstruction of a low-resolution map that shows an extra density in the DNA binding channel.

### Figure S8

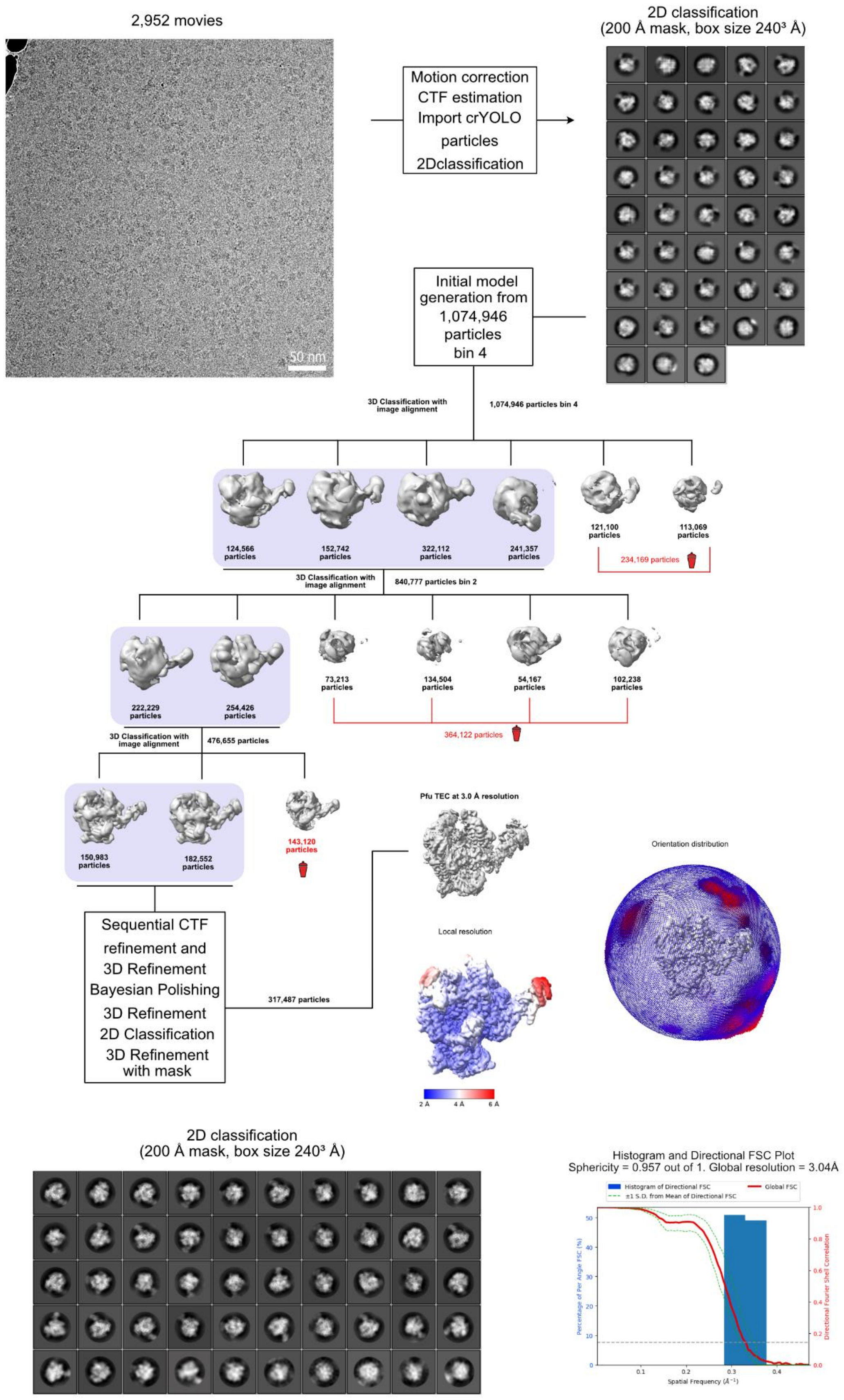
2,952 movies
50 nm
Motion correction
CTF estimation
Import crYOLO
particles
2Dclassification
2D classification
(200 Å mask, box size 240³ Å)
Initial model
generation from
1,074,946
particles
bin 4
3D Classification with
image alignment
1,074,946 particles bin 4
124,566
particles
152,742
particles
322,112
particles
241,357
particles
121,100
particles
113,069
particles
234,169 particles
3D Classification with
image alignment
840,777 particles bin 2
222,229
particles
254,426
particles
73,213
particles
134,504
particles
54,167
particles
102,238
particles
364,122 particles
3D Classification with
image alignment
476,655 particles
150,983
particles
182,552
particles
143,120
particles
Sequential CTF
refinement and
3D Refinement
Bayesian Polishing
3D Refinement
2D Classification
3D Refinement
with mask
317,487 particles
Pfu TEC at 3.0 Å resolution
Local resolution
2 Å
4 Å
6 Å
Orientation distribution
2D classification
(200 Å mask, box size 240³ Å)
Histogram and Directional FSC Plot
Sphericity = 0.957 out of 1. Global resolution = 3.04Å
Histogram of Directional FSC
±1 S.D. from Mean of Directional FSC
Global FSC
Percentage of Per Angle FSC (%)
Directional Fourier Shell Correlation
Spatial Frequency (Å⁻¹)

**Figure S8. Cryo-EM and single particle analysis processing workflow of the TEC sample including representative micrograph, 2D averages, 3D classifications, final refined map, local resolution heatmap, Fourier completeness plot, histogram and directional Fourier Shell Correlation (FSC).**

**Figure S9**

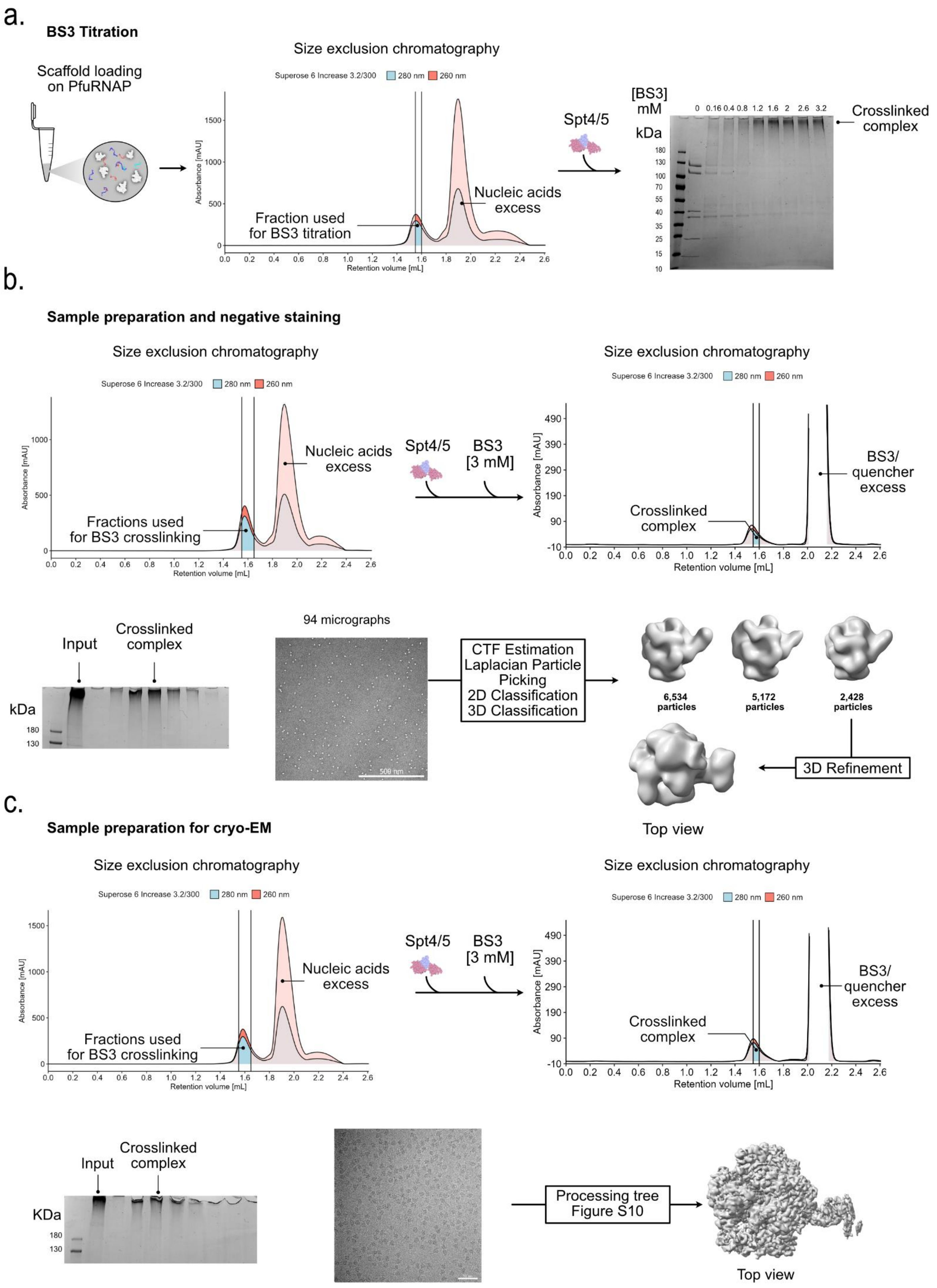
a.
BS3 Titration
Scaffold loading on PfuRNAP
Size exclusion chromatography
Superose 6 Increase 3.2/300 280 nm 260 nm
Absorbance [mAU]
Retention volume [mL]
Fraction used for BS3 titration
Nucleic acids excess
Spt4/5
[BS3] mM
0 0.16 0.4 0.8 1.2 1.6 2 2.6 3.2
kDa
180 130 100 70 55 40 35 25 15 10
Crosslinked complex
b.
Sample preparation and negative staining
Size exclusion chromatography
Superose 6 Increase 3.2/300 280 nm 260 nm
Fractions used for BS3 crosslinking
Nucleic acids excess
Spt4/5
BS3 [3 mM]
Crosslinked complex
BS3/ quencher excess
Input
Crosslinked complex
kDa
180
130
94 micrographs
500 nm
CTF Estimation
Laplacian Particle Picking
2D Classification
3D Classification
6,534 particles
5,172 particles
2,428 particles
3D Refinement
Top view
c.
Sample preparation for cryo-EM
Size exclusion chromatography
Superose 6 Increase 3.2/300 280 nm 260 nm
Fractions used for BS3 crosslinking
Nucleic acids excess
Spt4/5
BS3 [3 mM]
Crosslinked complex
BS3/ quencher excess
Input
Crosslinked complex
KDa
180
130
Processing tree Figure S10
Top view

**Figure S9. Sample preparation of TEC in complex with Spt4/5. a.** The TEC was formed in vitro by incubating *Pfu* RNAP with a synthetic elongation scaffold. The excess of nucleic acids was removed by size exclusion chromatography. The fraction containing the TEC with the highest concentration was incubated with recombinant Spt4/5 and crosslinked with different concentrations of BS3. Crosslinking success was verified using SDS PAGE analysis. **b.** The crosslinking reaction was optimized for EM purposes starting first with scaffold annealing to the RNAP, elimination of excess of nucleic acids (first chromatogram), crosslinking of Spt4/5 to the TEC using a final concentration of 3mM of BS3 with subsequent quenching the reaction. At this stage the excess of BS3, quencher and aggregates were eliminated by size exclusion chromatography (second chromatogram). The crosslinking success was assessed by SDS-PAGE. Negative staining was performed, and small dataset was acquired (96 micrographs), which was processed and yielded a low-resolution map showing an extra density on the clamp domain. **c.** The procedure described before for negative staining was repeated and cryo-grids were generated.

**Figure S10**

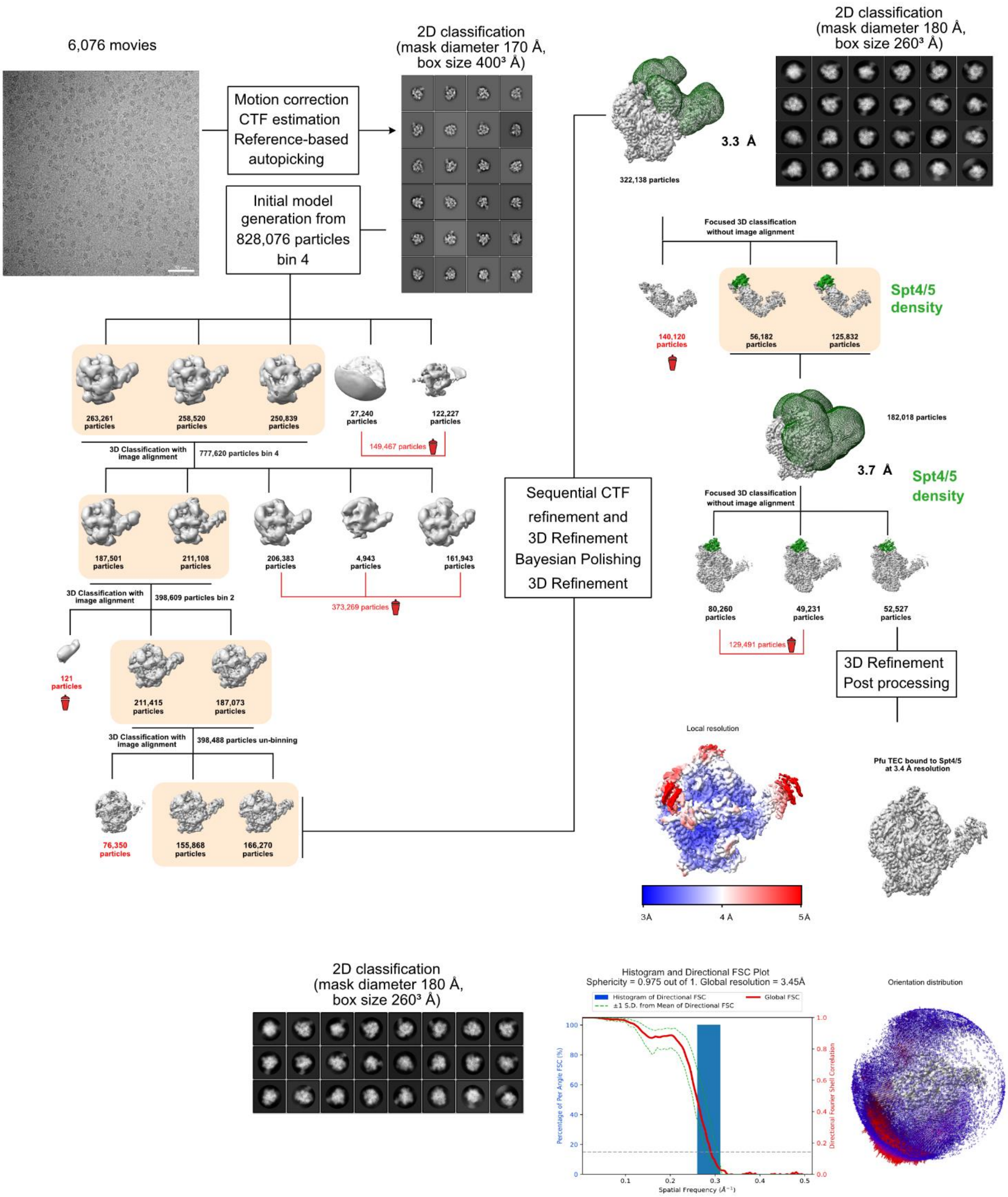


**Figure S10. Cryo-EM and single particle analysis processing workflow of the TEC in the presence of Spt4/5 (crosslinked sample) including representative micrograph, 2D averages, 3D classifications, final refined map, local resolution heatmap, Fourier completeness plot, histogram and directional Fourier Shell Correlation (FSC).**

**Figure S11**

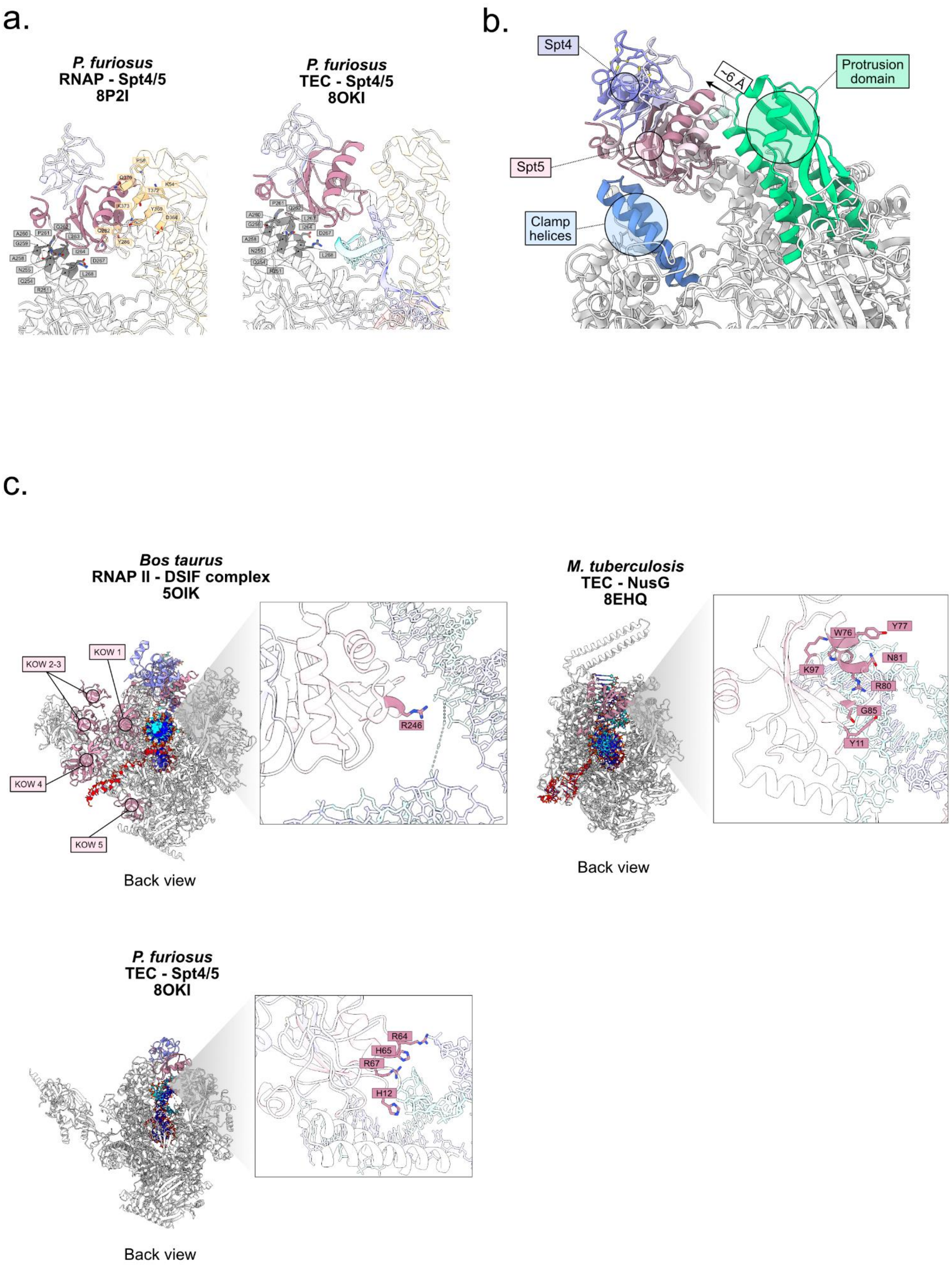

a.
P. furiosus
RNAP - Spt4/5
8P2I
P. furiosus
TEC - Spt4/5
8OKI
b.
Spt4
~6 Å
Protrusion domain
Spt5
Clamp helices
c.
Bos taurus
RNAP II - DSIF complex
5OIK
KOW 1
KOW 2-3
KOW 4
KOW 5
R246
Back view
M. tuberculosis
TEC - NusG
8EHQ
W76
Y77
K97
N81
R80
G85
Y11
Back view
P. furiosus
TEC - Spt4/5
8OKI
R64
H65
R67
H12
Back view

**Figure S11. Structural comparison of Spt4/5 binding behavior. a.** Comparison of the position and interactions of Spt4/5 interactions with *Pfu* RNAP. Interface analysis on ChimeraX 1.5 revealed the interaction pattern of Spt5 with RNAP in context of the TEC and in absence of nucleic acids. When the scaffold is absent, Spt4/5 is located ~6 Å further towards the DNA binding channel losing the interaction with the protrusion and lobe domain. **b.** Structural comparison of human RNAPII TEC in complex with Spt4/5 homolog, DSIF, the bacterial TEC with NusG from *Mycobacterium tuberculosis*, and the archaeal TEC bound to Spt4/5. **c.** Highlighted are amino acids of Spt5 or NusG that form interactions with the DNA in transcription elongation complexes. In eukaryotes (left) it has been described in Spt5-R246 is involved in transcriptional pausing[4]. Similarly, Vishwakarma et al. an interaction interface involving W76, Y77, R80, N81, T86, K97, Y11 and the non-template strand (middle) was described in Bacteria[5]. In this study, we investigated the effect of R64, H65, R67 and H12 on transcription elongation (right) in Archaea.

**Figure S12**

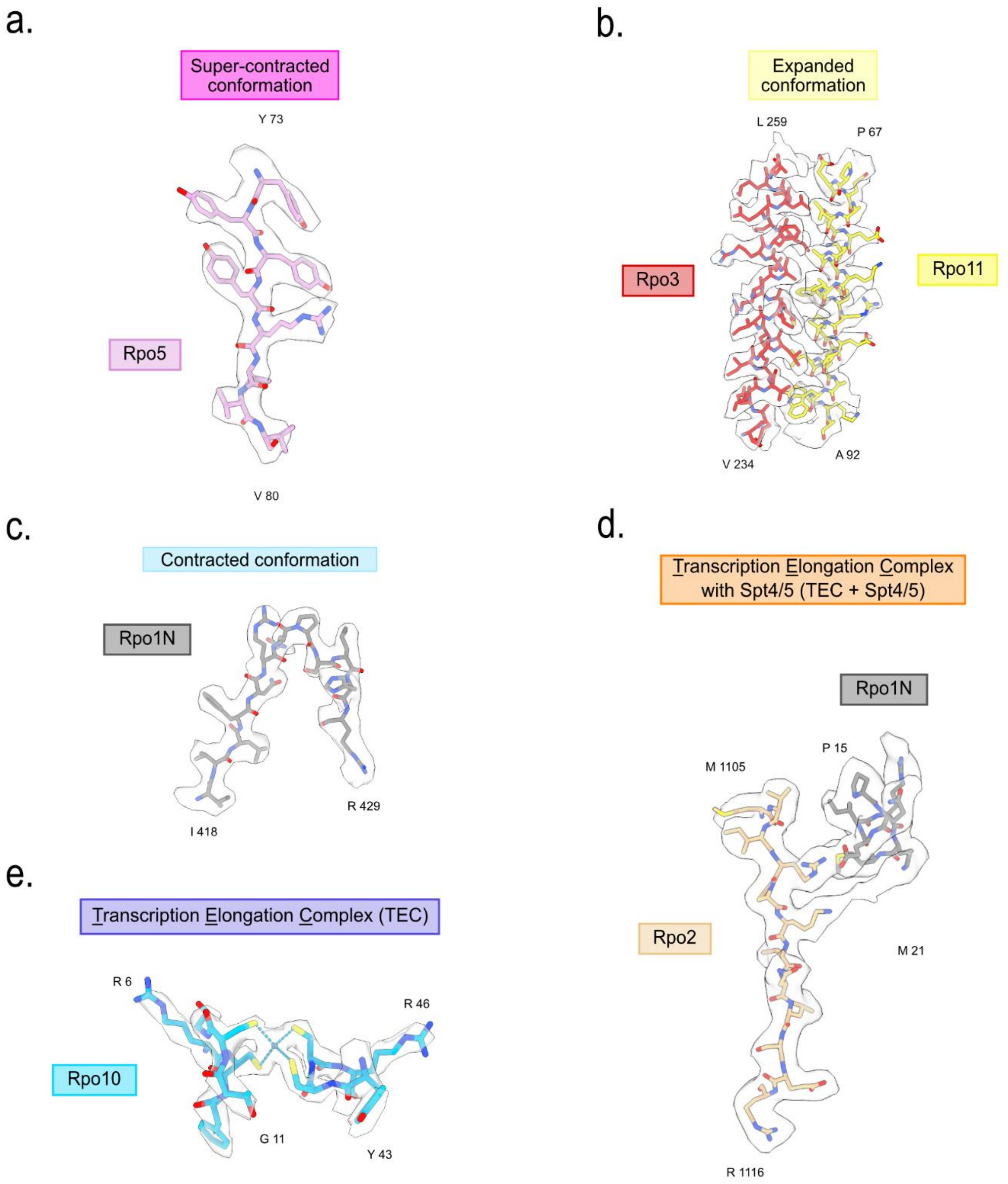


**Figure S12. Details of the different models and maps generated. a.** Rpo5 beta-sheet from residue $Y^{73}$ to $V^{80}$ with the respective density map corresponding to the *pfu* RNAP super-contracted conformation. **b.** Alpha-helices corresponding to Rpo3 ($V^{234}$-$L^{259}$) and Rpo11 ($P^{67}$-$A^{92}$) heterodimer with the respective density corresponding to the *pfu* RNAP expanded conformation. **c.** Detail of Rpo1N ($I^{418}$-$R^{429}$) with correspondent density from *pfu* RNAP contracted conformation. **d.** Detail of Rpo2 ($M^{1105}$-$R^{1116}$) and Rpo1N ($P^{15}$-$M^{21}$) and respective density map showing a continuous density linking the two lysins in the BS3 cross-linked TEC with Spt4/5 sample. **e.** Details of the zinc binding domain from subunit Rpo10 ($R^{6}$-$G^{11}$, $Y^{43}$-$R^{46}$) and respective density map from TEC sample.

**Figure S13**

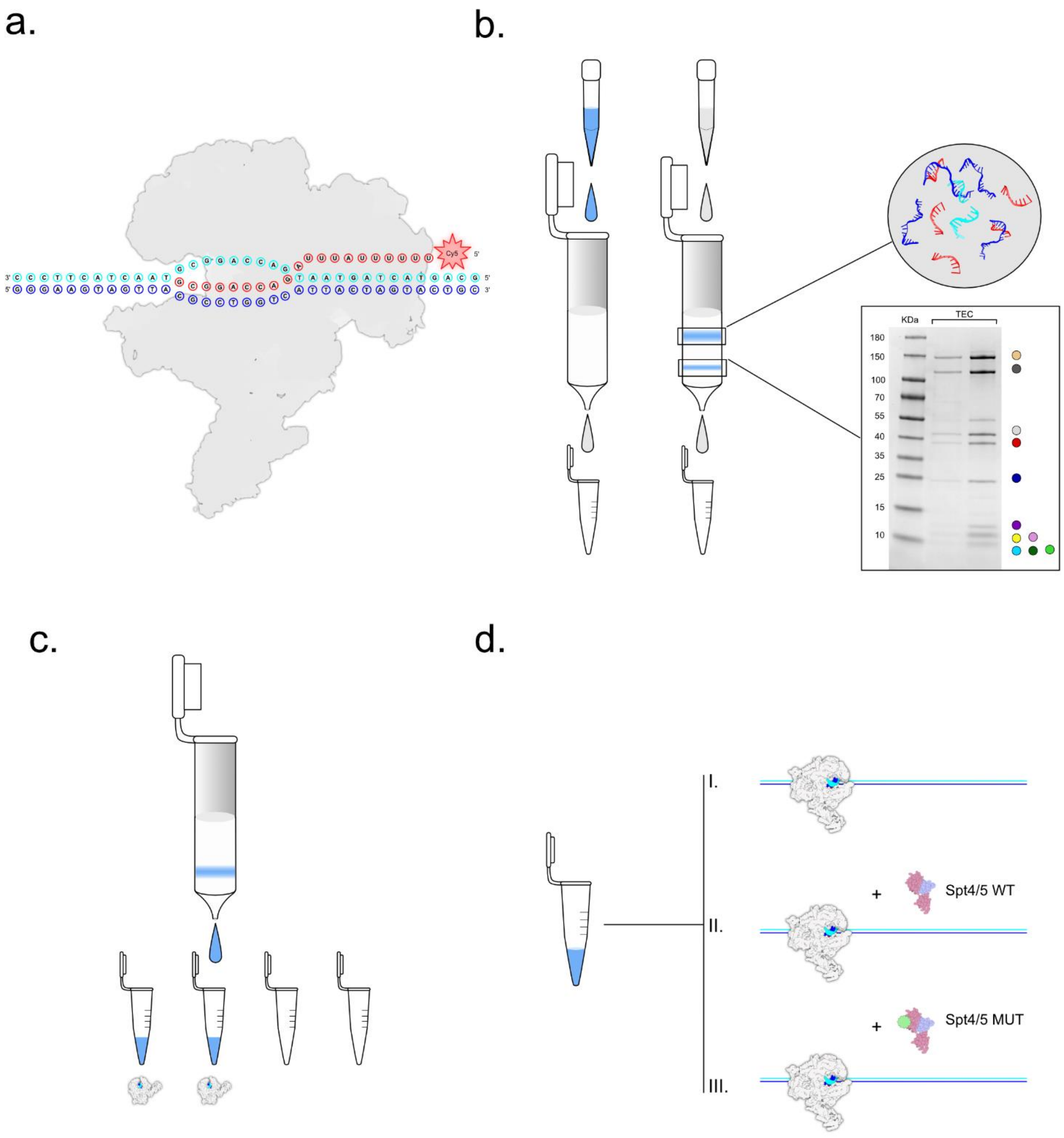


**Figure S13. TEC assembly for *in vitro* transcription assays. a.** *Pfu* RNAP was annealed to the synthetic scaffold containing a Cy5 labelled RNA (label is positioned at the 5'-end). **b.** The excess of nucleic acids was eliminated using gravity flow size exclusion chromatography columns. **c.** Fractions were manually collected and presence of the TEC was confirmed by SDS-PAGE as shown in panel b. **d.** *In vitro* transcription ivT) assays (ivT) were carried out using the following samples: i. ivT in absence of any co-factor, ii. ivT with additional Spt4/5 WT, iii. ivT mutated Spt4/5 variant HRHR-AAAA added.

**Table S1. Cryo-EM data collection and refinement statistics**

| | *Pfu* Apo RNAP Expanded conformation | *Pfu* RNAP with Spt4/5 | *Pfu* Apo RNAP Contracted conformation | *Pfu* Elongation complex | *Pfu* Elongation complex with Spt4/5 |
|---|---|---|---|---|---|
| **EMDB** | EMD-17130 | EMD-17366 | EMD-19033 | EMD-16809 | EMD-16929 |
| **PDB** | 8ORQ | 8P2I | 8RBO | 8CRO | 8OKI |
| **Data collection and processing** | | | | | |
| *Magnification* | 50k | 50k | 50k | 75k | 50k |
| *Voltage (kv)* | 200 | 200 | 200 | 300 | 200 |
| *Electron exposure (e⁻ /Å2)* | 40 | 40 | 40 | 88.73 | 40 |
| *Defocus range (μm)* | -1 to -2.2 | -1 to -2.2 | -1 to -2.2 | -1.2 to -2.0 | -1 to -2.2 |
| *Pixel size (Å)* | 0.968 | 0.968 | 0.968 | 1.0635 | 0.968 |
| *Symmetry imposed* | C1 | C1 | C1 | C1 | C1 |
| *Initial coordinates (no.)* | 1,209,249 | 1,209,249 | 1,183,000 | 1,229,339 | 1,878,102 |
| *Final particle (no.)* | 161,531 | 107,970 | 111,194 | 333,535 | 322,138 |
| *Map resolution (Å)* | 3.2 | 3.4 | 3.0 | 3.0 | 3.4 |
| *FSC threshold* | 0.143 | 0.143 | 0.143 | 0.143 | 0.143 |
| **Refinement** | | | | | |
| *Homology model used (PDB code)* | 6kf9 | 6kf9 | 6kf9 | 6kf9 | 6kf9 |
| *Model composition* | | | | | |
| *Non-hydrogen atoms* | 26195 | 27486 | 26375 | 27169 | 28561 |
| *Protein residues* | 3266 | 3430 | 3290 | 3289 | 3260 |
| *Nucleotides* | - | - | - | 39 | 52 |
| *Ligands* | 5(Zn), 1(Mg) | 6(Zn), 1(Mg) | 5(Zn), 1(Mg) | 5(Zn), 1(Mg) | 6(Zn), 1(Mg) |
| *APD (B factors)* | | | | | |
| *Protein (max)* | 98.61 | 245.70 | 98.61 | 214 | 245.60 |
| *Nucleotides (max)* | - | - | | 214 | 214.13 |
| *Ligand* | 30 | 30 | 30 | 30 | 30 |
| *Bonds (RMSD)* | | | | | |
| *Lengths (Å)* | 0.005 (0) | 0.004 (0) | 0.003 (0) | 0.003 (0) | 0.002 (0) |
| *Angles (°)* | 0.545 (0) | 0.626 (0) | 0.557 (11) | 0.548 (0) | 0.545 (5) |
| *Validation* | | | | | |
| *MolProbity score* | 1.69 | 1.57 | 1.53 | 1.50 | 1.28 |
| *Clashscore* | 8 | 8.44 | 9.3 | 10 | 5.62 |
| *Poor rotamers (%)* | 1.7% | 2% | 3% | 2.1% | 2% |
| *Ramachandran plot* | | | | | |
| *Favored (%)* | 96.54 | 97.41 | 97.83 | 98.28 | 98.09 |
| *Allowed (%)* | 3.46 | 2.59 | 2.17 | 1.72 | 1.88 |
| *Disallowed (%)* | 0 | 0 | 0 | 0 | 0 |

**Table S2. Table listing primers used to create the mutated *Pyrococcus furiosus* strain.**

| name | oligonucleotide sequence (5'→3') |
|---|---|
| 1648/1647-BHI-F | AGCAGCGGATCCGGCCCTTGAAAAG |
| PF1647_Fus-StrepII_Ri | CTTCTCGAACTGTGGGTGAGACCATGCGCTAGAGGTCAATTTTTGAAGTTCAC |
| Fus_Strep_His_F | CACCCACAGTTCGAGAAGCATCACCATCACCATCACTGAGGATGGGAGGTGTAAG |
| Fus_Strep_His_R | CTTACACCTCCCATCCTCAGTGATGGTGATGGTGATGCTTCTCGAACTGTGGGTG |
| PF1647upTk0914F | TGAGGATGGGAGGTGTAAGATATGAACTTTGAAGAGCTTGTTGAG |
| TK0914-R | CTAACGTCCAAGCTCCTTATGGGC |
| Tk0914PF1647dwF | GCCCATAAGGAGCTTGGACGTTAGTCCCTGCGGGGGT |
| Fus004_BamHI-R | CCACCTGGATCCTTTCCTCCCAG |


1. Jun, S.-H. *et al.* Direct binding of TFEα opens DNA binding cleft of RNA polymerase. *Nat Commun* **11**, 6123 (2020).

2. Pilotto, S. *et al.* Structural basis of RNA polymerase inhibition by viral and host factors. *Nat Commun* **12**, 5523 (2021).

3. Dienemann, C., Schwalb, B., Schilbach, S. & Cramer, P. Promoter Distortion and Opening in the RNA Polymerase II Cleft. *Molecular Cell* **73**, 97-106.e4 (2019).

4. Dollinger, R. *et al.* Assessment of the roles of Spt5-nucleic acid contacts in promoter proximal pausing of RNA polymerase II. *Journal of Biological Chemistry* **299**, 105106 (2023).

5. Vishwakarma, R. K., Qayyum, M. Z., Babitzke, P. & Murakami, K. S. Allosteric mechanism of transcription inhibition by NusG-dependent pausing of RNA polymerase. *Proc. Natl. Acad. Sci. U.S.A.* **120**, e2218516120 (2023).
